# ALTERATIONS IN PEROXISOMAL-MITOCHONDRIAL INTERPLAY IN SKELETAL MUSCLE ACCELERATES MUSCLE DYSFUNCTION

**DOI:** 10.1101/2024.04.25.591056

**Authors:** Marco Scalabrin, Eloisa Turco, Leonardo Nogara, Gaia Gherardi, Giulia Trani, Samuele Negro, Anais Franco Romero, Yorrick Jaspers, Elisa Baschiera, Rossella De Cegli, Eugenio Del Prete, Tito Cali, Bert Blaauw, Leonardo Salviati, Michela Rigoni, Cristina Mammucari, Sylvie Caspar-Bauguil, Cedric Moro, Marco Sandri, Stephan Kemp, Vanina Romanello

## Abstract

Whole-body energy expenditure, as well as glucose and lipid metabolism, are regulated by skeletal muscles, which account for 40-50% of human body mass. Peroxisomes are dynamic organelles that play a crucial role in lipid metabolism and clearance of reactive oxygen species, however their role in muscles remains poorly understood.

To clarify this issue, we generated a muscle-specific transgenic mouse line with peroxisome import deficiency resulting from deletion of peroxisomal biogenesis factor 5 (Pex5). Pex5 inhibition disrupted the tethering between peroxisomes and mitochondria, impaired lipid metabolism and reduced muscle force and exercise performance. Moreover, mitochondrial content and function were also altered, accelerating age-related structural defects, neuromuscular junction degeneration, and muscle atrophy. Altogether, our findings show the importance of preserving peroxisomal function and their contact sites with mitochondria to maintain muscle health during aging.

## INTRODUCTION

Skeletal muscle, the body’s largest tissue, plays a critical role in maintaining systemic metabolic balance through intricate interorgan communication. As a central hub for metabolic activities, it governs glucose and lipid balance and serves as the primary protein reservoir, supplying essential amino acids to fuel energy production in other organs during catabolic states. As such, precise adjustments in muscle mass and metabolic demands become indispensable for meeting overall metabolic needs and ensuring whole-body homeostasis ^1,2^. However, excessive catabolism during illness can exceed muscle plasticity, resulting in atrophy, and depleted metabolic reserves, leading to harmful functional limitations that impact disease onset and progression^1,2^.

Muscle atrophy and weakness represent significant clinical challenges observed in conditions such as cancer, diabetes, obesity, and cardiac failure, as well as unhealthy aging and infections like COVID-19. Muscle loss serves as a negative prognostic factor, causing respiratory insufficiency, loss of independence, and metabolic disruptions, thereby compromising life quality and elevating morbidity and mortality rates^1–3^. On the other hand, maintaining a healthy skeletal muscle mass is associated with a reduced risk of mortality^4,5^, underscoring the critical role of muscle health in overall body homeostasis.

Our understanding of pathways regulating muscle mass has greatly improved in the last years. However, the lack of effective therapeutic approaches for muscle wasting highlights our limited comprehension of the mechanistic insights involved in muscle atrophy. Molecular dissection of these mechanisms is crucial for paving the way toward successful drug development and intervention strategies.

Peroxisomes are ubiquitous dynamic metabolic organelles, adjusting their number and protein content to cellular metabolic needs. In mammals, they harbor over 50 anabolic and catabolic matrix enzymes involved in essential metabolic pathways, such as fatty acid oxidation, biosynthesis of plasmalogen and bile acids, and reactive oxygen species detoxification^6^. As non-autonomous organelles, peroxisomes closely interact with other organelles^6^, particularly mitochondria, through physical and functional connections like membrane contact sites (MCS), mitochondrial-derived vesicles, and biological messengers like ROS or lipids^6–9^. Thus, in light of the close interaction between the two organelles, it is not surprising that the functional impairment of either organelle is likely to induce dysfunction to the other^10^. Despite the clear interplay between peroxisomes and mitochondria, their specific contributions to pathology are not fully understood^8^. While the impact of mitochondrial metabolic activity on muscle function has been extensively explored^11^, peroxisomes in skeletal muscle have been poorly investigated. The regulation of peroxisomes and their potential contribution to muscle function remain unknown, representing a critical gap in our understanding, particularly considering the significant metabolic role of these organelles.

Peroxisome biogenesis relies on the coordinated activity of several peroxins (Pex) proteins to assemble and maintain functional peroxisomes. Mutations in at least 14 different Pex genes result in rare autosomal recessive Peroxisomal Biogenesis Disorders (PBD), also known as Zellweger Spectrum Disorders ^12,13^. The inability to form functional peroxisomes in PBD leads to the loss of essential peroxisomal metabolic functions and subsequent multisystem tissue pathology. PBD are a heterogeneous group of disorders ranging from severe to relatively milder phenotypes, with severity inversely related to age of onset. The most severe presentation is lethal within the first year of life, characterized by craniofacial dysmorphism, neuronal dysfunction, hepatorenal failure, and profound muscular hypotonia^12^.

Biochemically, PBD are characterized by the accumulation of very-long-chain (VLCFA) and branched-chain fatty acids, bile acid intermediates, pipecolic acid, and severe depletion of plasmalogens and docosahexaenoic acid^12^.

In line with the intense metabolic activity of skeletal muscle, peroxisome absence or dysfunction severely impacts muscle tissue in PBD patients. Muscle biopsies from individuals with mutations in Pex12 and Pex16 reveal a secondary mitochondrial myopathy characterized by enlarged mitochondria, reduced mitochondrial respiratory chain activity, lipid accumulation, and muscle atrophy. These pathological features likely contribute to clinical manifestations such as generalized hypotonia, respiratory issues, and sucking difficulties^14,15^. Pex5, an essential receptor protein crucial for importing most peroxisomal enzymes into the peroxisomal lumen, holds particular significance. In humans, mutations in the Pex5 gene result in PBD^16^, and the total deletion of Pex5 in mice recapitulates PBD, resulting in early postnatal mortality^17^. Importantly, the diaphragm in these mice is the most severely affected muscle, exhibiting several mitochondrial abnormalities. This observation suggests that respiratory failure may, in part, contribute to the mortality observed in some patients^18^.

Moreover, peroxisomal dysfunction is associated with aging, and age-related diseases such as diabetes, obesity, cancer, and neurodegenerative disorders^19,20^. Notably, these conditions share muscle atrophy and dysregulated muscle function as common features. However, the role of peroxisomes in muscle function remains largely unexplored, and the physiological significance of peroxisomal-mitochondrial cooperation in muscle health and disease remains unclear.

In this study, we show that muscle-specific deletion of the peroxisomal biogenesis factor Pex5 in mice triggers early changes in lipid and amino acid metabolism. In addition, it reduces the tethering between peroxisomes and mitochondria, resulting in a detrimental effect on muscle force and exercise performance. These disruptions progressively contribute to a decline in mitochondrial content and function, together with sarcomere and neuromuscular junction degeneration, accumulation of aggregates, which altogether lead to muscle atrophy and induce the premature onset of muscle aging.

## RESULTS

### MUSCLE-SPECIFIC ABLATION OF PEX5 RESULTS IN THE IMPAIRMENT OF PEROXISOME ASSEMBLY, PROTEIN IMPORT, AND PEXOPHAGY FLUX

The physiological role of peroxisomes in skeletal muscle remains significantly underexplored. To investigate their relevance in maintaining skeletal muscle metabolism and mass, we generated a mouse model to induce peroxisomal dysfunction specifically within the muscle tissue. We crossed Pex5 floxed mice^21^ with a transgenic line expressing Cre recombinase (CRE) under the control of the Myosin Light Chain 1 fast promoter (MLC1f promoter)^22^, thereby generating mice lacking Pex5 in skeletal muscle from birth (MLC-Pex5 ^-/-^). The deletion of Pex5 in muscle tissue was validated through Real-Time PCR (Fig.1A) and western blot analyses (Fig.1B) in tibialis anterior (TA), demonstrating a significant reduction in both Pex5 transcript and protein levels. Muscle-specific Pex5^-/-^ knockout animals (hereafter referred to as KO) were born at the expected Mendelian ratio and exhibited full viability, fertility, and physical appearance indistinguishable from their Pex5^fl/fl^ littermates (hereafter referred to as “Control”). Accordingly, the postnatal changes in body weight gain, and the lean and fat mass body composition were similar in both control and KO groups (Suppl. Fig.1A-C).

**Figure 1.**
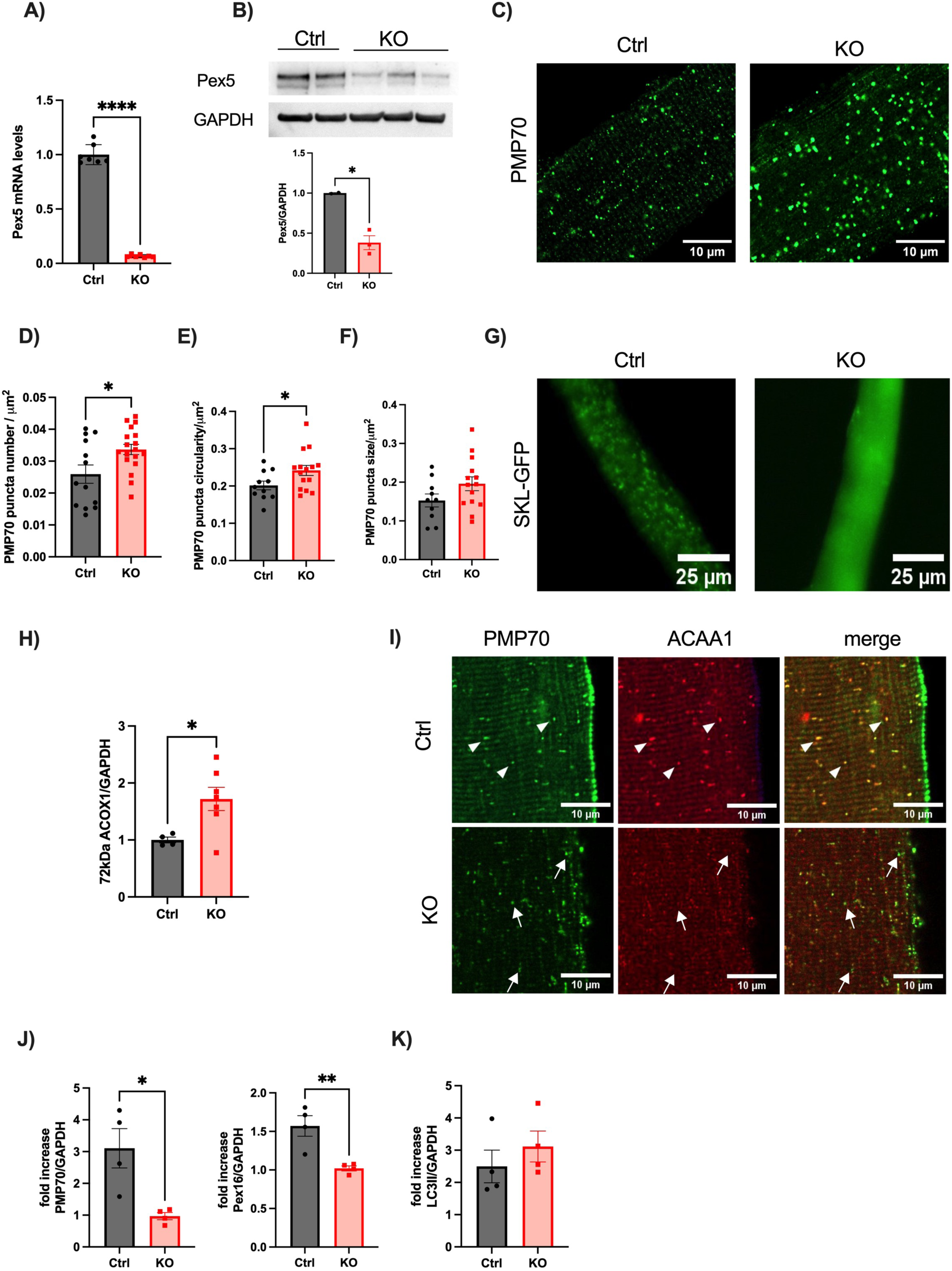
Pex5 deletion in skeletal muscle impairs peroxisome assembly, protein import and pexophagy flux. Pex5 mRNA (A) and Pex5 protein levels (B) are downregulated in tibialis anterior muscle of KO mice. GAPDH, glyceraldehyde-3-phosphatedehydrogenase. C) PMP70 immunostaining in isolated FDB muscle fibers. Quantification of PMP70-positive puncta number (D), circularity (E), and size (F) was normalized to fiber area. G) Peroxisomal import capacity was investigated by transfecting SKL-GFP into FDB fibers. Confocal microscopy analysis shows GFP-positive puncta in Control (Ctrl) fibers, while KO fibers display GFP cytosolic distribution, indicating peroxisomal protein import impairment in KO muscle fibers. H) Densitometric analysis of western blots of the 72kDa cytosolic ACOX1 band protein levels (ACOX1 antibody PT 10957-1-AP). I) Representative confocal microscopy images showing PMP70 and ACAA1 immunostaining in FDB fibers. Arrowheads indicate PMP70-positive structures containing ACAA1 in control fibers, while arrows show PMP70-positive structures in KO muscle with reduced ACAA1 signaling compared to control. J) Fold increase of PMP70, Pex16. K) Fold increase of LC3II. Western blot of total muscle homogenates probed with PMP70, Pex16, and LC3 were normalized to GAPDH and plotted as a ratio between colchicine treated samples and paired samples without colchicine. Data are presented as mean ± SEM (with individual data points). Unpaired two-tailed Student’s t tests (A,B,D,E,F,H,J,K) were used. Statistical significance: *p ≤ 0.05; **p ≤ 0.01; ***p ≤ 0.001.

To visualize muscle peroxisomes, we performed an immunostaining against the peroxisomal membrane protein PMP70 (ABCD3), a well-established marker of peroxisomes, in isolated flexor digitorum brevis (FDB) muscle fibers at 3 months of age (Fig.1C). Consistent with findings in fibroblasts from patients with Pex5 mutations^23^, we observed an increase in the number of PMP70-positive puncta (Fig.1D) and a more circular morphology, indicative of reduced elongation, in KO fibers compared to the control group (Fig.1E). Additionally, peroxisomal size displayed a trend of increase in KO fibers, although this trend did not reach statistical significance (Fig.1F).

SKL is the peroxisomal targeting signal 1 (PTS1) signal recognized by Pex5 to import the peroxisomal proteins from the cytosol to the peroxisomal matrix. To investigate the protein import capacity of PMP70-positive structures in the KO muscle, we conducted in vivo transfection of SKL-GFP plasmids into FDB fibers of 3-month-old mice. The control fibers displayed a punctate distribution, while KO fibers exhibited a cytosolic distribution (Fig.1G), reflecting an impairment in the peroxisomal protein matrix import. Using immunoblot analysis we investigated the peroxisomal import capacity by characterizing the acyl-CoA oxidase-1 (ACOX1) enzyme, which represents the first and rate-limiting enzyme in peroxisomal fatty acid oxidation. ACOX1 is synthesized as a 72 kDa precursor, which, upon import into the peroxisome, undergoes proteolytic processing to yield 52kDa and 20kDa forms^24^. Consistent with peroxisomal dysfunction, the 72 kDa ACOX1 full length protein accumulates in KO fibers (Fig.1H and Suppl.Fig.1D). This accumulation suggests a disruption in the peroxisomal import and processing pathways that normally lead to the formation of the 52 kDa and 20 kDa forms. However, our attempts to detect the 52 kDa and 20 kDa processed bands by western blot analyses, using two different antibodies in control and KO skeletal muscle fibers yielded no discernible results (Suppl.Fig.1D). This outcome suggests a potential limitation either in the specificity of the antibodies employed or in the tissue characteristics. The acetyl-CoA Acyltransferase 1 (ACAA1) protein plays a crucial role in the third phase of very-long-chain fatty acid (VLCFA) β-oxidation^25^. Initially synthesized in the cytosol, ACAA1 is subsequently imported into the peroxisomal matrix. Immunohistochemistry indicated reduced co-localization between ACAA1 and PMP70 in KO muscle samples (Fig.1I). This observation confirms that the residual PMP70-positive peroxisomal structures in KO muscle are import deficient. Consequently, PMP70-positive structures in KO fibers resemble peroxisomal ghosts, characterized by import-deficient residual peroxisomal membranes resulting from aberrant peroxisome assembly, with little or no matrix content. Importantly, such structures are well-documented hallmarks of PBD patients with Pex5 mutations ^23,26^ as well as a distinctive feature of the total Pex5 KO animal model ^17^.

PEX5 has been reported as a target for ubiquitination that promotes in mammals, the autophagic degradation of peroxisomes, known as pexophagy^27–29^. Therefore, to further explore the dynamics of PMP70-positive structures, we investigated the turnover of peroxisomes though the autophagic pathway in vivo by administering vehicle or colchicine to 3 months old mice. Colchicine is known to block the fusion of autophagosomes with lysosomes^30^. After colchicine treatment, we investigated pexophagy flux in the tibialis anterior muscle by quantifying the levels of peroxisomal membrane proteins PMP70 and Pex16 using Western Blot analysis (Fig.1J and Suppl. Fig.1E). Additionally, we quantified LC3 II to assess general autophagy in accordance with the latest autophagy guidelines^30^ (Fig.1K and Suppl. Fig.1E). Importantly, Pex5 deletion reduces the turnover of PMP70 and Pex16 without affecting the general autophagy flux, in line with previous observations^27^.

In summary, our findings suggest that the decreased pexophagy flux in the absence of Pex5 likely contributes to the accumulation of import-deficient peroxisomal ghosts in the KO muscle tissue. Moreover, our data emphasize the muscle-specific Pex5 KO mouse as a suitable model for unraveling the consequences of peroxisomal dysfunction in skeletal muscle.

### PEX5 DELETION IN SKELETAL MUSCLE INDUCES EARLY ALTERATIONS IN LIPID METABOLISM

Peroxisomes perform crucial roles in lipid metabolism, including fatty acid oxidation of very long chain fatty acids (VLCFA) and ether lipids biosynthesis such as plasmalogens. To gain deeper insights into peroxisomal function, we performed untargeted lipidomics on gastrocnemius muscle (GNM) from control and KO mice at an early (3 months) and late time points (18 months). This approach allowed us to track the dynamic changes in lipid metabolism over time. We identified more than 2000 lipids covering multiple lipid classes. Muscle tissue from Pex5-deleted mice exhibited significant alterations in their lipid profiles at both 3 and 18 months-old, as shown by principal component analysis (PCA) (Suppl.Fig.2A). Consistent with the disruption of peroxisomal lipid metabolism, KO muscles showed a reduction in ether lipids and increased levels of VLCFA containing lipid species at all ages (Fig.2A and Suppl.Fig.2B). Specifically, the total levels of plasmalogen species, phosphatidylcholine (PC[P]) and phosphatidylethanolamine (PE[P]), as well ether-linked phosphatidylcholine (PC[O]) and phosphatidylethanolamine (PE[O]), were already diminished in KO muscle tissue at the 3-month time point (Fig.2B and Suppl. Fig.2C). Furthermore, we observed increased levels of phosphatidylcholine (PC), phosphatidylethanolamine (PE), and triglycerides (TG) containing VLCFA with a tendency toward longer unsaturated fatty acid (FA) chains (Fig.2C). More specifically, PC and PE species exhibited a shift towards longer acyl chain lengths, while species with acyl chains shorter than C40 were reduced (indicated by a dashed line in Fig.2C). However, the total levels of PC, PE, diacylglycerol (DG), and TG remained unaltered in KO muscle compared to control muscle across all age groups, with exceptions noted for PE, cholesteryl ester (CE), and sphingomyelin (SM), which exhibited increases at 18 months (Suppl.Fig.2D). Quantitative targeted lipidomic analyses on 9 months old muscle confirmed no change in total phospholipids, DG and TG between genotypes also at this age (Supp. Fig.2E). While total ceramide levels (Cer d) showed no significant differences between control and KO at both timepoints (Suppl.Fig.2D), specific species within the ceramides lipid class, such as Cer d31:0, Cer d41.0, Cer d43.2, exhibited significant increases in muscles from 18 months KO mice (Suppl. Fig.2F). Additionally, Cer d38.1 was consistently induced in both KO muscles at 3 and 18 months (Suppl.Fig.2F).

**Figure 2.**
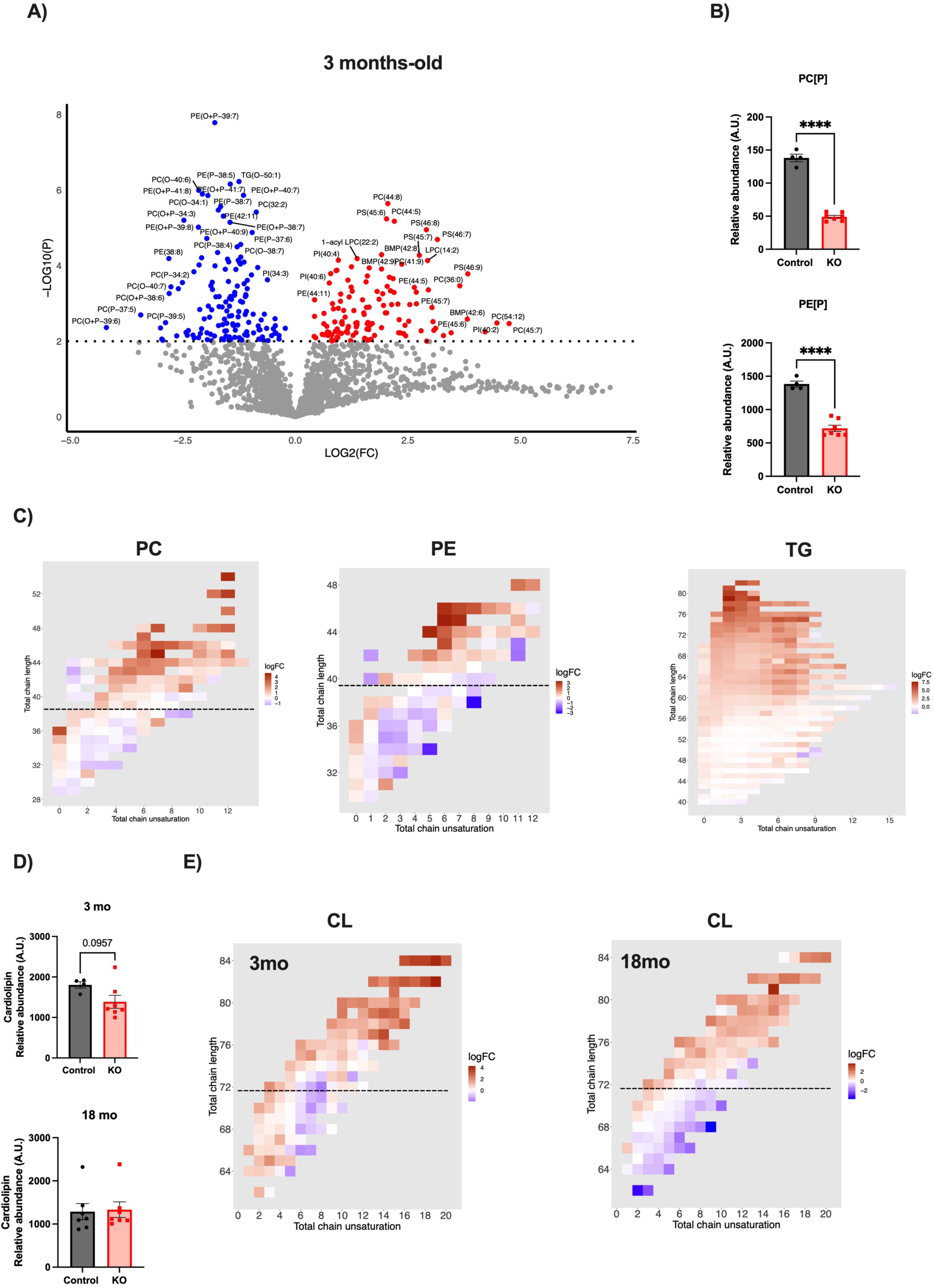
Alterations in Pex5-null muscle lipidomic profile. A) Volcano plot based on untargeted lipidomics analysis performed in 3 months-old gastrocnemius muscles comparing control vs. KO. Significant decreases are depicted in blue, while significant increases are depicted in red. B) Skeletal muscle concentration of total phosphatidylcholine (PC[P]), and phosphatidyletanolamine (PE[P]) plasmalogen species is reduced in KO muscle. C) Phosphatidylcholine (PC), phosphatidylethanolamine (PE) and triglycerides (TG) with longer unsaturated acyl chains increase in 3 months-old KO muscle. The dashed line indicates C40 acyl chains. D)Total cardiolipin (CL) muscle concentration in skeletal muscle at 3 and 18 months-old. CL chain length over C72 and unsaturation increases in KO muscle at both ages. The dashed line indicates C72. All lipidomics data were normalized to dry tissue weight. Data are presented as mean ± SEM (with individual data points). Welch t-test was used in (A). Unpaired two-tailed Student’s t tests or Mann-Whitney test were used for experiments comparing two groups (B, C, and D). Statistical significance: *p ≤ 0.05; **p ≤ 0.01; ***p ≤ 0.001.

Cardiolipin (CL) is a phospholipid synthesized in the inner mitochondrial membrane, crucial for maintaining proper cristae folding, respiratory chain integrity, and ATP synthase function^31^. We observed a non-significant trend towards a reduction of total levels of cardiolipin (CL) at 3 months-old time point and no changes at 18 months-old (Fig.2D). However, the composition of CL species was altered across all age groups, showing a tendency toward very long unsaturated fatty acid (FA) chains, while acyl chains below C72 were reduced (indicated by a dashed line in Fig.2E), in line with PBD patient’s fibroblasts lipidomic profile^32^.

### MITOCHONDRIAL CONTENT UNDERGOES AGE-DEPENDENT DOWNREGULATION IN PEX5-DEFICIENT MUSCLE

To investigate the network of genes controlled by Pex5, we used an unbiased approach, performing bulk RNA sequencing (RNA-seq) analysis on the gastrocnemius muscle of 3-, 9 and 18-months-old and age-matched control littermates.

Despite substantial alterations in the lipidomic profile of KO muscle already at 3 months (Supp.Fig.2A), the most significant metabolic changes emerged in the muscle transcriptome at 9 and 18 months. We performed both Gene Ontology Enrichment Analysis (GOEA) within the “biological process” and cellular components categories and Gene set enrichment analysis (GSEA) restricting the output to biological processes (BP) and cell components (CC) and KEGG pathway gene sets. Moreover, a custom GSEA was performed to verify the significant modulation of the mitochondrial respiration process. These analyses consistently associated Pex5 deletion with the downregulation of genes involved in mitochondrial respiration, mitochondrial ATP synthesis, fatty acid oxidation, and the generation of precursor metabolites and energy (Fig. 3A, 3B, Suppl.Fig.3A, and Suppl. Table 3A, 3B). Accordingly, a cluster of 25 Differentially Expressed Genes (DEGs) closely associated with mitochondrial functions, particularly mitochondrial respiration, and fatty acid oxidation, exhibited downregulation exclusively at the 9- and 18-month timepoints (Fig.3C, and Suppl. Table 3C). Consistent with a reduction of mitochondrial transcripts, the protein levels of the mitochondrial outer membrane protein VDAC (Fig.3D and Suppl. Fig.3B) are reduced in isolated mitochondria from KO muscle at 9 and 18 months. In addition, the subunits of mitochondrial respiratory complexes in KO muscle-isolated mitochondria were reduced at 9 and 18 months, while at 3 months, they resembled controls, except for a trend of increase of the subunit MTCO1 of complex IV (Fig.3E and Suppl. Fig. 3B). Of note, given the observed reduction in several mitochondrial protein levels (not shown), VDAC and OXPHOS subunits protein levels were normalized to total protein content. Importantly, the age-dependent decline in mitochondrial transcripts and proteins in KO muscle is associated with progressive changes in mitochondrial DNA (mtDNA) (Fig.3F) and citrate synthase activity (Fig.3G), two commonly used markers of mitochondrial content^33,34^.

**Figure 3.**
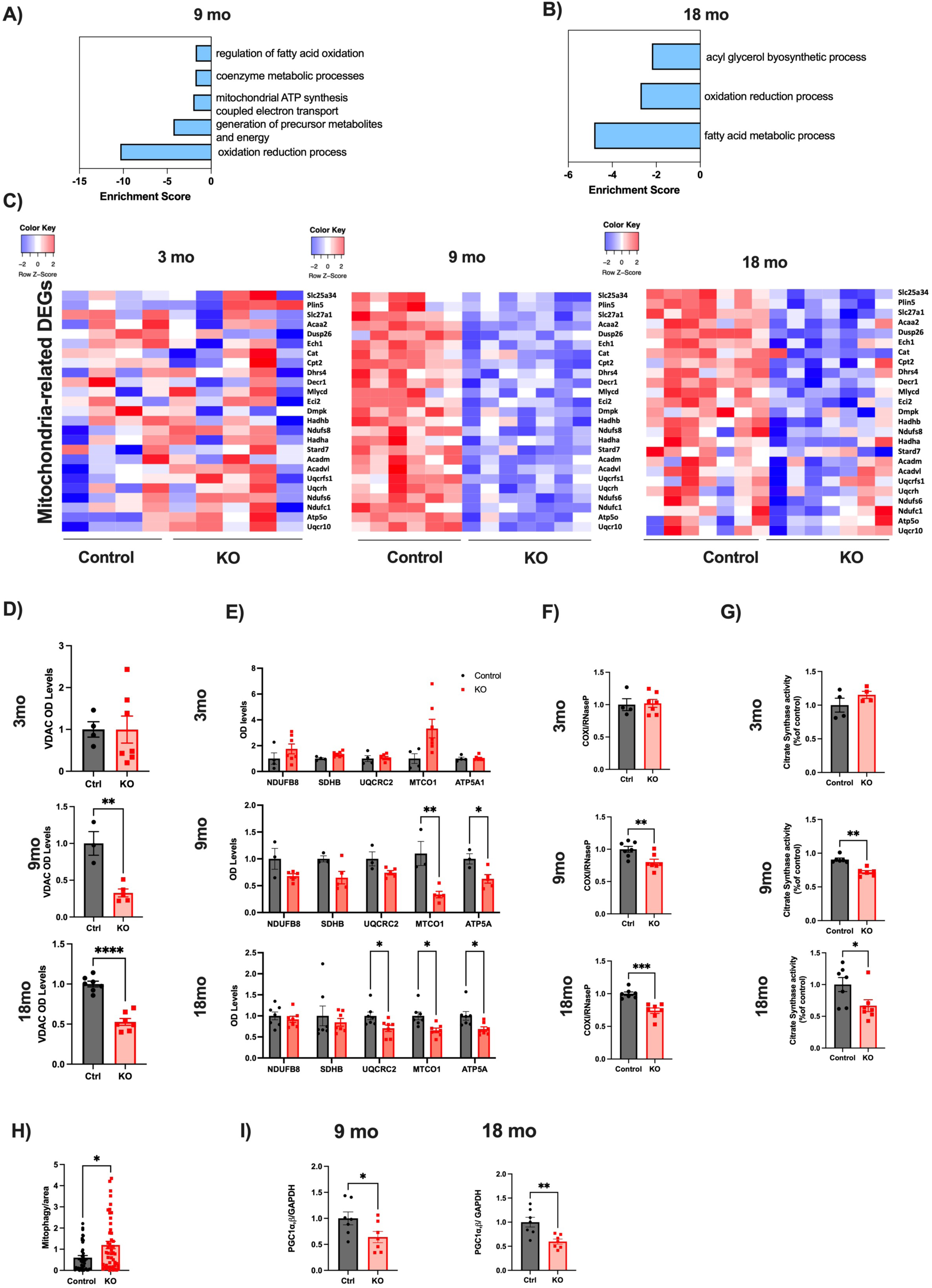
Ablation of Pex5 in skeletal muscle results in progressive mitochondrial content decline. (A-B) Transcriptomic RNA-seq Gene Ontology Enrichment Analysis (GOEA) analysis showing the downregulation of genes involved in metabolic pathways. A) Biological processes (BP) significantly inhibited at 9 months of age. The GO analysis was performed restricting the output to the BP terms in which the 58 inhibited genes at 9 months of age are mainly involved. The Enrichment Score of each BP term is plotted (refer to Suppl. Table Fig 3A). B) BP significantly inhibited at 18 months of age. The GO analysis was performed restricting the output to the BP terms in which the 48 inhibited genes at 18 months of age are mainly involved. The Enrichment Score of each BP term is plotted (refer to Suppl. Table Fig 3B). C) Heatmap on 25 mitochondria-related genes significantly inhibited in KO vs CTR at 9 months of age (refer to Suppl. Table Fig 3C). The heatmap show the expression of these transcripts at 3, 9 and 18 months of age. DEGs: Differentially Expressed Genes. (D-E) Densitometric quantification of immunoblots performed with isolated mitochondria from control and KO muscle normalized by total protein content at different ages showing VDAC (D) and mitochondrial respiratory complexes subunits protein levels (E). Mitochondrial DNA copy number (F) and citrate synthase activity quantification (G) were measured in TA muscle at 3,9, and 18 months-old time points. Data are normalized to controls. H) Mitophagy flux was analyzed by electroporation of a reporter plasmid (mt-mKEIMA) into flexor digitorum brevis muscles of 9 months-old control and KO mice. Changes in fluorescent spectra were detected and normalized to fiber area. I) Densitometric quantification of PGC1α/β by western blot, normalized to GAPDH protein levels, was performed in muscles at 9 and 18 months-old. Data are presented as mean ± SEM (with individual data points). Unpaired two-tailed Student’s t tests (D,F,H,I), Multiple unpaired two-tailed Student’s t tests (E) or Mann-Whitney test (G) were used for experiments comparing two groups. Statistical significance: *p ≤ 0.05; **p ≤ 0.01; ***p ≤ 0.001.

Mitochondrial content depends on the balance between mitochondrial degradation and mitochondrial biogenesis. To analyze if the reduction in mitochondrial content is explained by the degradation of damaged and dysfunctional mitochondria through selective autophagy, known as mitophagy, we employed mitochondrial-targeted Keima probe (mt-Keima) transfection to assess the mitophagy flux in FDB fibers. We observed an induction of mitophagy flux in KO muscle fibers at 9 months (Fig.3H) along with a progressive reduction in the protein levels of the master regulators of mitochondrial biogenesis PGC1α and PGC1β at both 9 and 18 months, thereby resulting in decreased mitochondrial content at this age (Fig.3I, and Suppl. Fig.3C).

Overall, these data indicate that Pex5 ablation in skeletal muscle results in a gradual decrease in mitochondrial content due to increased turnover of mitochondria by mitophagy, which is not counteracted by mitochondrial biogenesis. This likely exerts a substantial impact on metabolic signatures over time.

### DISRUPTION OF PEROXISOMAL-MITOCHONDRIAL INTERPLAY PRECEDES ALTERATIONS IN MITOCHONDRIAL ULTRASTRUCTURE AND FUNCTION IN PEX5-NULL MUSCLES

Peroxisomes and mitochondria work in synergy, and physical connections between these organelles at membrane contact sites are critical for efficient metabolic intermediate transfer to achieve key metabolic processes. Our data regarding changes in Pex5-null muscles in peroxisomal function, lipid metabolism, and mitochondrial content point to alterations in the peroxisomal mitochondrial metabolic interaction. However, it remains unclear whether these changes are a consequence of physical contact disruption between both organelles in the absence of Pex5. To explore whether peroxisome-mitochondria membrane contacts sites (MCS) are indeed remodeled in Pex5-KO muscle fibers, we overexpressed in vivo in FDB muscle fibers the split-GFP-based contact site sensor (SPLICS)^35^, capable of detecting interactions occurring within a range of 8-10nm. The reporter is engineered to express equimolar amounts of the organelle-targeted non-fluorescent GFP β-strand 11 and the GFP1-10 portions of the superfolded GFP protein variant in a single vector. These two portions of the GFP can reconstitute fluorescence when two opposing membranes come into proximity^36^. The peroxisomal-mitochondrial SPLICS reporter contains the C-terminal domain of the human ACBD5 protein as a peroxisomal targeting sequence in the β-strand 11 and the outer mitochondrial membrane Tom20 N33 targeting sequence in the GFP1–10 moiety^37^. Importantly, the inhibition of Pex5 is sufficient to reduce peroxisome-mitochondria contact sites in KO muscle fibers already at 3 months (Fig.4A and Suppl. Fig. 4A). The early disruption of the physical contact between both organelles in the KO muscle could lead to progressive metabolic alterations affecting mitochondrial ultrastructure and function. Accordingly, analysis of mitochondrial ultrastructure in the extensor digitorum longus (EDL) muscle through electron microscopy reveals significant changes over time. At 3 months, KO muscle closely resembles that of the control group, with an electron-dense matrix, parallel internal cristae, and mitochondria positioned at the I band near the Z lines (Fig.4B). However, by 9 months, some sporadic mitochondria in KO muscle exhibit swelling and disrupted cristae structures (Fig.4B). In contrast, at 18 months, the mitochondrial ultrastructure reverts to a state like the 3-month condition (Fig.4B). This is paralleled with a progressive increase in mitochondrial size in KO muscle from 3 to 18 months (Fig. 4C).

**Figure 4.**
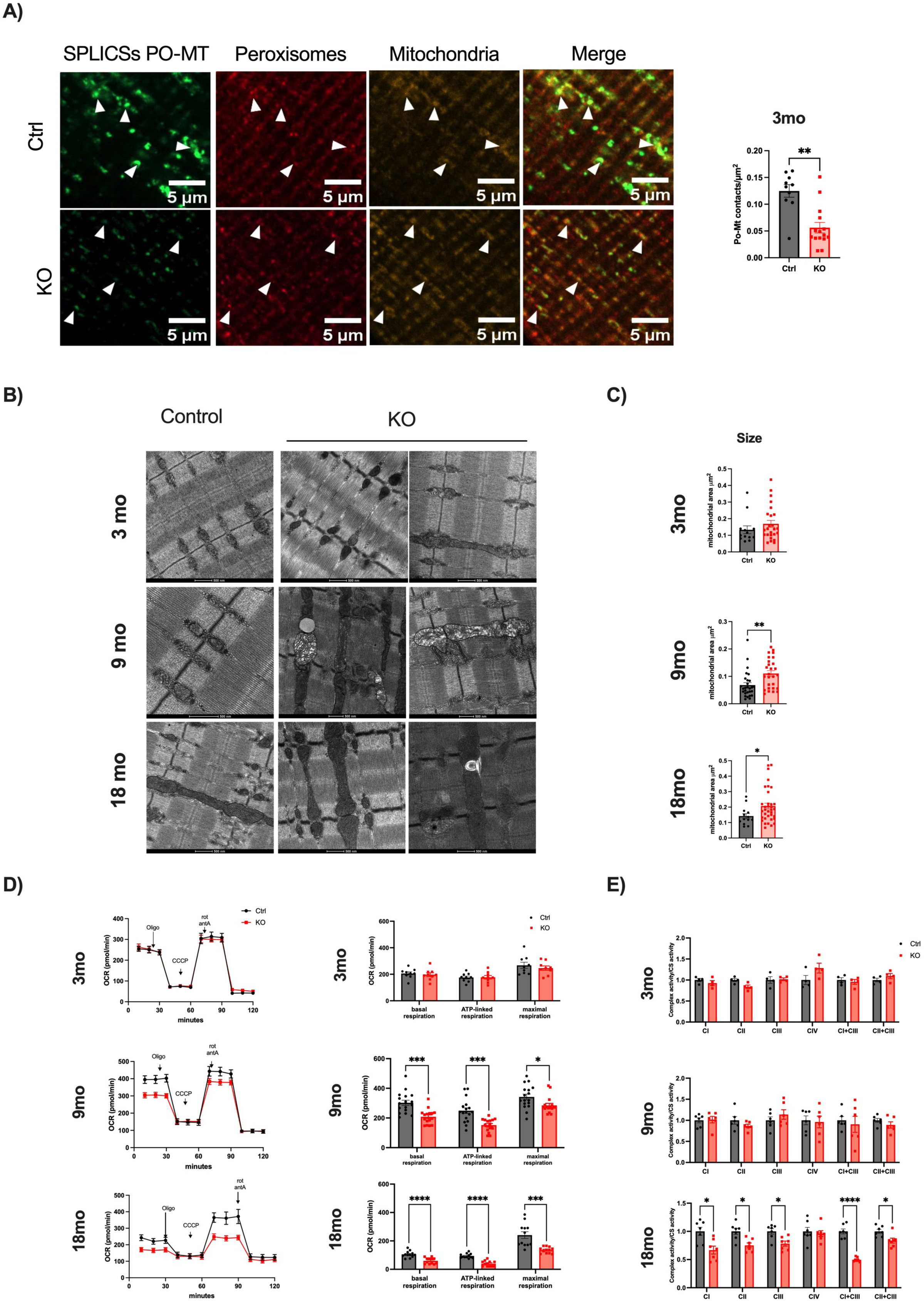
Impact of Pex5 inhibition on peroxisomal-mitochondrial-contact sites, mitochondrial ultrastructure, size, and function. A) Left: Higher magnification of representative images displayed in Suppl. Fig. 4E showing FDB fibers from 3 months-old mice expressing the Po/Mt SPLICS_s_-P2A^PO-MT^ probes. These images allow the identification of the co-localization of GFP (SPLICSsPO-MT) fluorescent dots with the targeted organelles, specifically peroxisomes labeled with anti-PMP70 antibody, and mitochondria labeled with anti-Tom20. Right: quantification of the Po/Mt SPLICS-P2A (short) contacts. The SPLICS dots were quantified from the 3D rendering of a complete Z-stack. B) Representative electron micrographs of EDL muscles of controls and KO mice. C) Quantification of mitochondrial area measurement in EM images of EDL muscle. D) OCR measurements indicated age-dependent reduced respiratory capacity in KO FDB myofibers compared to controls. Left: representative traces. Right: quantification. To calculate basal and maximal respiration, non-mitochondrial O2 consumption was subtracted from absolute values. ATP-linked respiration was calculated as the difference between basal and oligomycin-insensitive O2 consumption. Data are normalized on mean calcein fluorescence. E) Respiratory complex single enzyme activity normalized to citrate synthase activity. Data are presented as mean ± SEM (with individual data points). Unpaired two-tailed Student’s t test (A), Mann-Whitney test (C), and multiple unpaired two-tailed Student’s t test (D and E) were used for experiments comparing two groups. Statistical significance: *p ≤ 0.05; **p ≤ 0.01; ***p ≤ 0.001.

In line with these morphological changes, the oxygen consumption rate (OCR), normalized by the fluorescence of total fiber calcein protein content, remained unchanged at 3 months but showed a gradual decline in KO muscles from 9 to 18 months. This decline affected the basal, ATP-linked, and maximal respiration (Fig.4D). Additionally, measurement of the electron transport chain (ETC) mitochondrial respiratory complexes activity showed an age-dependent decrease. Normalization of the complex activities by citrate synthase activity, a marker of mitochondrial content, showed significant changes only at 18 months (Fig.4E). This indicates that the reduction in mitochondrial content, already evident at 9 months, precedes the onset of mitochondrial dysfunction at 18 months.

In addition, a set of DEGs related to fatty acid catabolism, displayed downregulation specifically at 9 and 18 months, with no such changes observed at 3 months (Suppl.Fig.4B and Suppl. Table 4B). Supporting these findings, real-time PCR showed the downregulation of genes associated with mitochondrial β oxidation in KO muscles. Specifically, ACADM was downregulated at 9 and 18 months, while CPT1, CPT2, and ACADL were downregulated at 18 months, with no significant changes observed at 3 months. These results suggest potential alterations in mitochondrial lipid catabolism (Suppl. Fig.4C).

Collectively, these results suggest that the downregulation of Pex5 leads to the disruption of the physical connection between peroxisomes and mitochondria, progressively affecting metabolite exchange between the two organelles. Over time, this disturbance likely impacts mitochondrial content, ultrastructure, and overall function.

### PEX5 DELETION IN MUSCLE RESULTS IN EARLY MUSCLE DYSFUNCTION

To explore the physiological role of Pex5 in controlling skeletal muscle mass, we focused on our mRNA sequencing data analysis, which revealed a clear association between Pex5 deletion, and the downregulation of transcripts associated with myofibril organization, muscle contraction, and muscle structure development at 3 months of age (Fig.5A, and Suppl. Table 5A). Notably, twelve genes involved in these processes exhibited significant downregulation (Fig. 5B, and Suppl. Table 5B). Additionally, untargeted metabolomics analysis revealed significant differences in the intramuscular levels of amino acids in 3 months-old KO mice. Interestingly, KO muscle exhibited increased levels of glutamine, glutamate, asparagine, valine, aspartate, histidine, tryptophan, phenylalanine, and phosphoserine compared to controls (Fig.5C and Suppl. Fig.5A). The accumulation of these amino acids has been associated with enhanced protein breakdown in conditions characterized by metabolic dysregulation, muscle weakness, and muscle loss^38,39^. To investigate if the transcriptomic and metabolic alterations correlate with functional defects, we assessed hindlimb muscle force in live 3-month-old animals subjected to contractions after electrical stimulation at increasing frequencies until tetanus was reached. The force frequency curve (Fig. 5D) and maximal tetanic force (100Hz) (Fig.5E), normalized for gastrocnemius muscle mass (Suppl. Fig. 5B), revealed reductions at 3 months and 18 months. Notably, muscle weakness was not associated with muscle mass loss, as cross-sectional area measurements of TA fibers at 3 months indicated no differences in size distribution between control and KO fibers (Fig.5F). However, by 9 months, there was a significant reduction in the number of fibers, with cross-sectional areas ranging from 2500 to 3000 μm² (Fig.5G). By 18 months, KO muscle fibers showed an increased prevalence of smaller sizes, ranging from 1000 to 1500 μm², accompanied by a decrease in the number of fibers ranging from 3000 to 4000 μm² (Fig. 5H). The alterations in fiber size were unrelated to changes in fiber type distribution, which remained consistent across all ages in both control and KO muscles (Suppl. Fig.5C).

**Figure 5.**
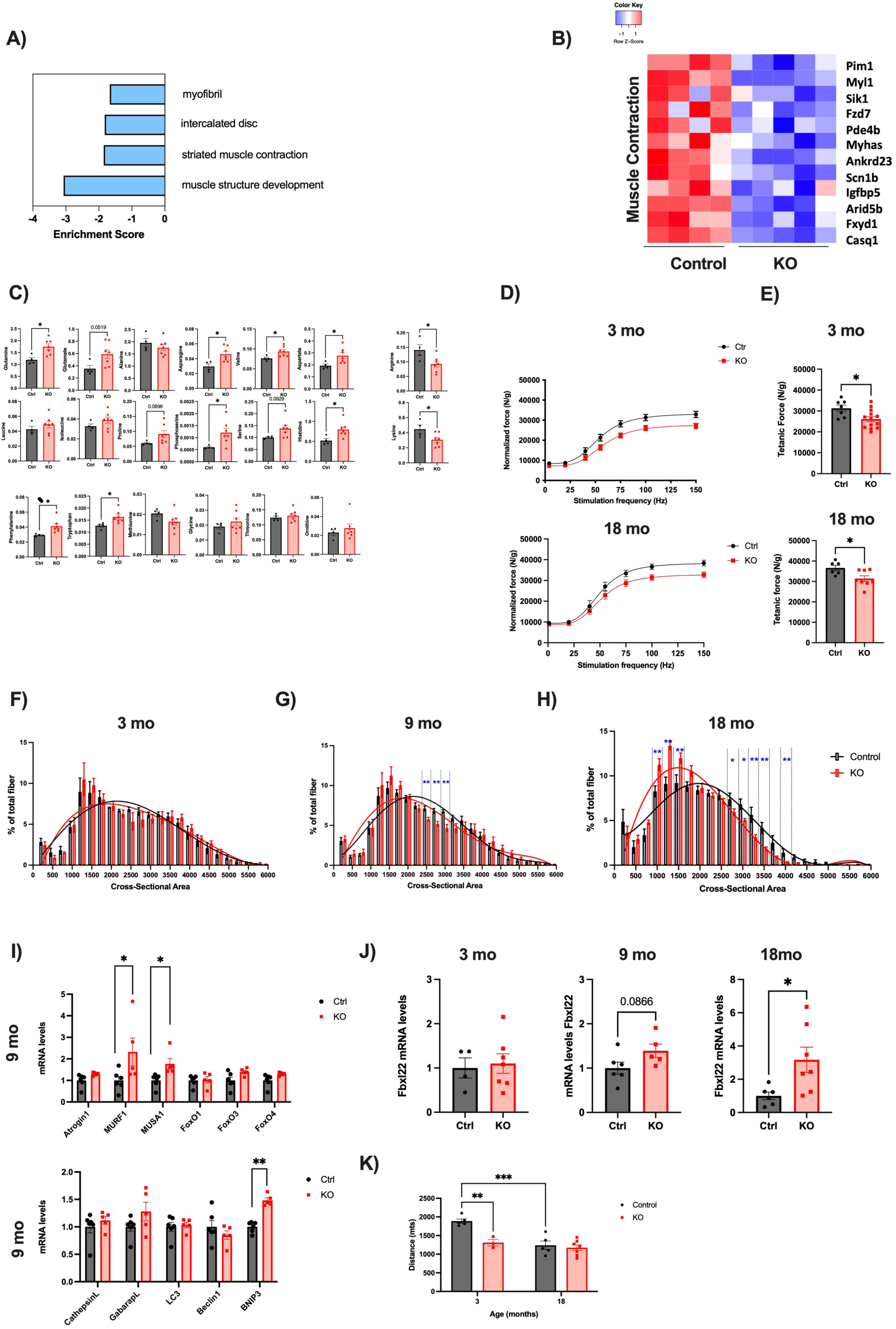
Pex5 deletion in skeletal muscle results in early-onset muscle weakness, which precedes muscle atrophy. A) Biological Processes (BP) and Cellular Components (CC) significantly inhibited at 3 months of age. The GO analysis was performed restricting the output to the BP and CC terms in which the 30 inhibited genes at 3 months of age are mainly involved. The Enrichment Score of each term is plotted (refer to Suppl. Table Fig 5A). B) Heatmap on 12 muscle contraction-related genes significantly inhibited in KO vs CTR GNM muscle at 3 months of age (refer to Suppl. Table Fig. 5B). C) Amino acids concentration in GNM muscle from 3 months old mice. All metabolomics data were normalized to dry tissue weight. D) Normalized force frequency curve of control and KO mice at 3 and 18 months of age. Force measurements were performed in vivo on gastrocnemius muscles and normalized to muscle wet weight. E) Normalized maximal tetanic force obtained at 100 Hz stimulation. F-H) Fiber size distribution of control and KO TA muscle at 3 (F), 9 (G), and 18 months of age (H). I) mRNA levels of atrophy-related genes in TA muscle of control and KO mice at 9 months. Upper panel: ubiquitin-proteasome-related genes. Lower panel: autophagy-related genes. J) Fbxl22 mRNA levels in TA muscles at 3, 9, and 18 months. K) Running distance on treadmill until exhaustion of control and KO mice at 3 and 18 months of age. Data are presented as mean ± SEM (with individual data points). Unpaired two-tailed Student’s t tests (C, E, J) or multiple unpaired two-tailed Student’s t test (F,G,H,and I) were used for experiments comparing two groups. A two-way ANOVA with Sidak’s multiple comparisons test (E) was performed for experiments comparing more than two groups (K). Statistical significance: *p ≤ 0.05; **p ≤ 0.01; ***p ≤ 0.001.

To explore the signaling pathways that contribute to muscle atrophy, we analyzed the levels of atrogenes, which are genes commonly altered in catabolic conditions^40^. At 9 months, the ubiquitin ligases MuRF1 and MUSA1 were upregulated in the KO muscle (Fig.5I). Additionally, the expression of the mitophagy gene Bnip3 increased in the KO muscle, consistent with the enhanced mitophagy flux observed at this age (Fig.3H). However, by 18 months, mRNA levels of atrogenes related to both the ubiquitin-proteasome and autophagy systems resembled those in the control muscle, except for FoxO1, which showed upregulation in the KO muscle (Suppl. Fig.5D).

This shift from induction to normalization of atrogenes between 9 and 18 months suggests that the activation of these genes precedes the onset of muscle loss in the KO mice. In addition, our RNA-seq data highlighted the induction of the novel F-box E3 ubiquitin ligase Fbxl22 in KO muscle at 18-month-old (Suppl. Fig.5E). Prior studies have demonstrated that Fbxl22 overexpression is sufficient to induce muscle atrophy ^41^. Consistent with the gradual changes observed in the cross-sectional area of KO muscles (Fig.5F-H), real-time PCR analysis showed the increase of Fbxl22 mRNA levels from 9 to 18 months in Pex5-deleted muscle (Fig.5J).

To further evaluate the consequences of Pex5 ablation on muscle performance in vivo, we subjected mice to treadmill exercise until exhaustion. Interestingly, 3-month-old KO mice covered approximately 30% less distance before reaching exhaustion compared to controls (Fig.5K). This suggests that Pex5 plays a crucial role in sustaining endurance during exercise, a role that becomes apparent even at this early stage, coinciding with initial changes in lipid metabolism. It’s noteworthy that the reduced exercise capacity observed in 3-month-old KO mice mirrors the age-related decline seen in both control and KO mice at 18 months (Fig.5K).

Collectively, these findings indicate that the alterations in transcriptomic, lipidomic, and metabolic profiles triggered by peroxisomal dysfunction in the skeletal muscle of 3-month-old KO mice trigger premature muscle weakness, preceding the progressive muscle atrophy observed between 9 and 18 months. Moreover, the similarity in running capacity between young KO mice and 18-month-old control mice suggests an accelerated decline in exercise endurance associated with Pex5 deficiency, positioning young knockout mice at a performance level like that of older control mice.

### AGE-ASSOCIATED MYOPATHY AND NEUROMUSCULAR JUNCTION DEGENERATION OCCURS EARLIER IN PEX5 KO MICE

To assess the impact of Pex5 deletion on muscle health, we performed a histological analysis of the TA muscle through H&E staining in both control and KO mice. At 3 months, the H&E staining revealed no signs of muscle degeneration, regeneration, or inflammation in KO muscle sections (Fig. 6A). However, at both 9 and 18 months, KO muscle exhibited a similar increase in the number of center-nucleated muscle fibers compared to age-matched controls (Fig. 6A). The mispositioning of myonuclei is a common hallmark of myofiber degeneration and regeneration observed in human myopathies and aging sarcopenia^42^.

**Figure 6.**
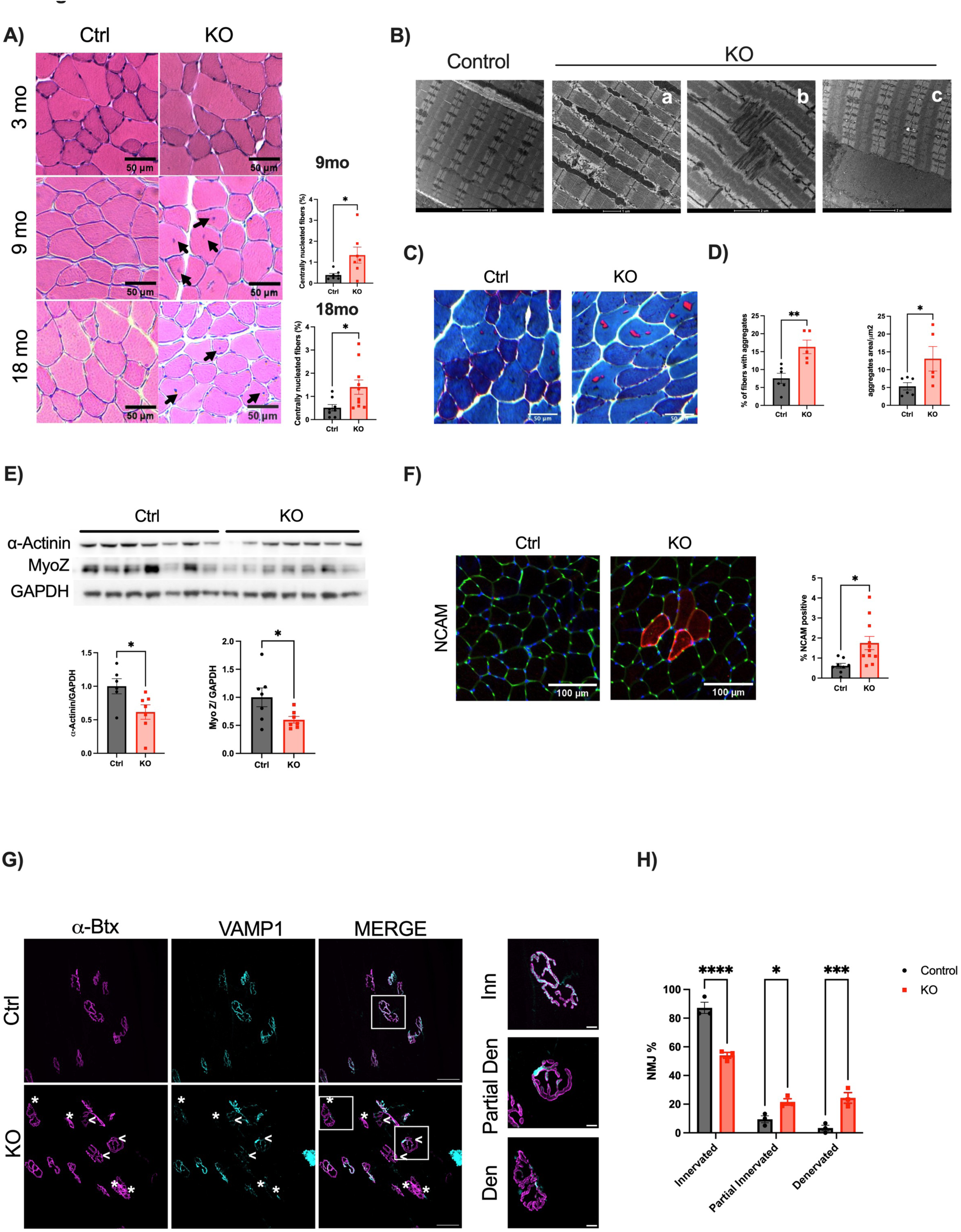
Pex5-null muscles exhibit an early development of the sarcopenic features. *A)* Left: representative Hematoxylin-Eosin staining of TA. The arrows indicate center nuclei in both 9- and 18-months KO mice. Right. Quantification of the number of fibers with center nuclei relative to total fiber number in the muscle section. B) Representative images of electron micrographs of EDL muscles from control and KO mice showing ultrastructural defects in 18 months-old KO muscle. C) Representative images of modified Gomori trichrome staining of TA muscle from control and KO mice at 18 months of age. D) Left: Quantification of the number of fibers with tubular aggregates (in red) relative to the total fiber number in the muscle section. Right: Quantification of the aggregate area normalized to the fiber area. E) Upper panel: total protein extracts from TA muscles were immunoblotted with the indicated antibodies. Lower panel: Densitometric analysis. Data are normalized to GAPDH protein levels. F) Left: Representative image showing NCAM positive fibers in TA KO muscles at 18 months. Right: Quantification of the number of NCAM-positive fibers relative to the total number of fibers in the muscle section. G) EDL muscles were processed for indirect immunofluorescence using fluorescent α-BTx to stain post-synaptic AChRs (magenta), and anti-VAMP1 antibodies to identify the pre-synaptic compartment (cyan). Asterisks identify denervated NMJs while arrows partial denervated ones. Scale bar: 50 µm. Magnification of innervated, partial denervated and denervated NMJs is shown in the right panel (10 µm). H) Quantification of innervated, partial denervated, and denervated NMJs in EDL muscles. N = 3, 30 NMJs analyzed/muscle. Data are presented as mean ± SEM (with individual data points). Unpaired two-tailed Student’s t tests (A, D, E, F) were used for experiments comparing two groups and a two-way ANOVA with Sidák’s multiple comparisons test (E). Statistical significance: *p ≤ 0.05; **p ≤ 0.01; ***p ≤ 0.001.

Therefore, to investigate myofiber integrity, we monitored the muscle ultrastructure of the EDL muscle by electron microscopy. We observed a consistent accumulation of ultrastructure defects over time. At 3 months, muscle ultrastructure in KO is like controls, exhibiting a regular sarcomeric structure (Suppl. Fig.6A, upper panel). By 9 months, KO muscles presented sporadic disorganized sarcomere arrangements (Suppl. Fig.6A, lower panel). However, at 18 months KO muscles exhibited a higher frequency of sarcomere alterations (Fig.6Ba), including Z-line smearing (Fig.6Bb), and the presence of tubular aggregates (Fig.6Bc). Tubular aggregates, derive from sarcoplasmic reticulum (SR) expansion and are associated with several skeletal muscle disorders, as well as normal and accelerated aging^43,44^. In light of this, we use modified Gömöri trichrome staining to examine 18 months-old muscles (Fig.6C). We quantified both the number of fibers presenting the tubular aggregates and the area occupied by these aggregates within each fiber, which appeared as bright red subsarcolemmal inclusions within the tibialis anterior sections. KO muscles show a significant increase in the percentage of fibers displaying tubular aggregates, with a corresponding increase in the area occupied within the fibers (Fig.6D).

To better understand why sarcomere integrity is affected by Pex5 deletion we focused on the novel F-box E3 ubiquitin ligase Fbxl22, which is upregulated in KO muscle at 18 months old (Fig.5J and Suppl Fig.5E). Fbxl22 has been reported to promote sarcomeric turnover by targeting Z-line proteins, including filamin-C and α-actinin, for proteasomal degradation in cardiomyocytes^45^, while in skeletal muscle, only α-actinin has been demonstrated as an Fbxl22 target substrate for ubiquitination so far^41^. Immunoblot analysis of Z-line proteins revealed a substantial reduction of approximately 40% in α-actinin and 35% in MyoZ protein levels within the KO muscle at 18-months (Fig.6E), while filamin-C and CapZ protein levels remained unchanged (Suppl. Fig.6B). Therefore, Fbxl22 induction likely leads to progressive disassembly of the sarcomere structure, potentially contributing to the observed muscle atrophy in the KO muscle (Fig.5F-H). Moreover, Fbxl22 is early induced during muscle denervation, and its knockdown partially protects against denervation-induced muscle loss^41^. Because FoxO1 gene was upregulated in knockout mice and FoxOs play a critical role in atrogenes regulation^46^, we hypothesized that also Fbxl22 is under FoxO regulation. Notably, several putative FoxO-binding elements in the promoter region of Fbxl22 have been identified by bioinformatic analysis. To investigate whether FoxOs control Fbxl22 expression, we used a muscle-specific mouse model lacking FoxO1, FoxO3, and FoxO4, which we have previously characterized^40^. Our findings further support this correlation by revealing a significantly upregulation of Fbxl22 in the muscles of control mice after 3 days of denervation. Conversely, Fbxl22 upregulation was blunted in denervated FoxOs-deficient mice, suggesting Fbxl22’s dependence on FoxO signaling (Suppl. Fig.6C). Since Fbxl22 is upregulated during denervation, and we observed increased Fbxl22 transcripts levels in 18 months-old KO muscle (Fig.5J and Suppl Fig.5E), we investigated whether loss of myofiber innervation could occur in Pex5 deficient muscle. Real time PCR analysis at 18 months showed the significant upregulation of established denervation markers such as muscle-associated receptor tyrosine kinase (MUSK) and runt-related transcription factor 1 (RUNX) ^47,48^ (Suppl. Fig.6D). The expression of the neural cell adhesion molecule (NCAM), which is typically enriched in the postsynaptic endplates of the neuromuscular junction (NMJ) and transiently redistributed on muscle fibers upon denervation, was significantly increased in the 18-month KO muscle (Fig.6F). Next, we analyzed the NMJ morphology by immunostaining the presynaptic side for the synaptic vesicle protein VAMP1 and the post-synaptic NMJ component with fluorescently labeled α-bungarotoxin, which tightly binds to acetylcholine receptors (AChRs). Innervated fibers were identified by the complete overlap observed by confocal microscopy between the pre- and post-synaptic components. Partial denervation and complete denervation were recognized by the partial or complete mismatch between the two signals. EDL muscle from control mice exhibited an almost complete colocalization of synaptic vesicle VAMP1 and postsynaptic AChRs. Conversely, 18 months-old KO mice exhibited a variable degree of mismatch between pre- and post-synaptic structures, with an increased proportion of partial and complete denervated NMJs and a reduction of the innervated NMJs compared to control EDL muscle (Fig. 6G, H).

In conclusion, peroxisome assembly deficiency in skeletal muscle disrupts the interaction between peroxisomes and mitochondria, triggering metabolic alterations in lipid and amino acid homeostasis, and negatively affecting muscle force and exercise performance. Gradually, these changes first impact mitochondrial content, which subsequently impairs mitochondrial function. Consequently, these cumulative effects results in muscle atrophy, and the development of a myopathic phenotype. This phenotype is characterized by the accumulation of tubular aggregates, proteolytic breakdown of sarcomeres, and degeneration of the neuromuscular junction, hence contributing to the accelerated development of aging sarcopenia (Fig.7).

**Figure 7.**
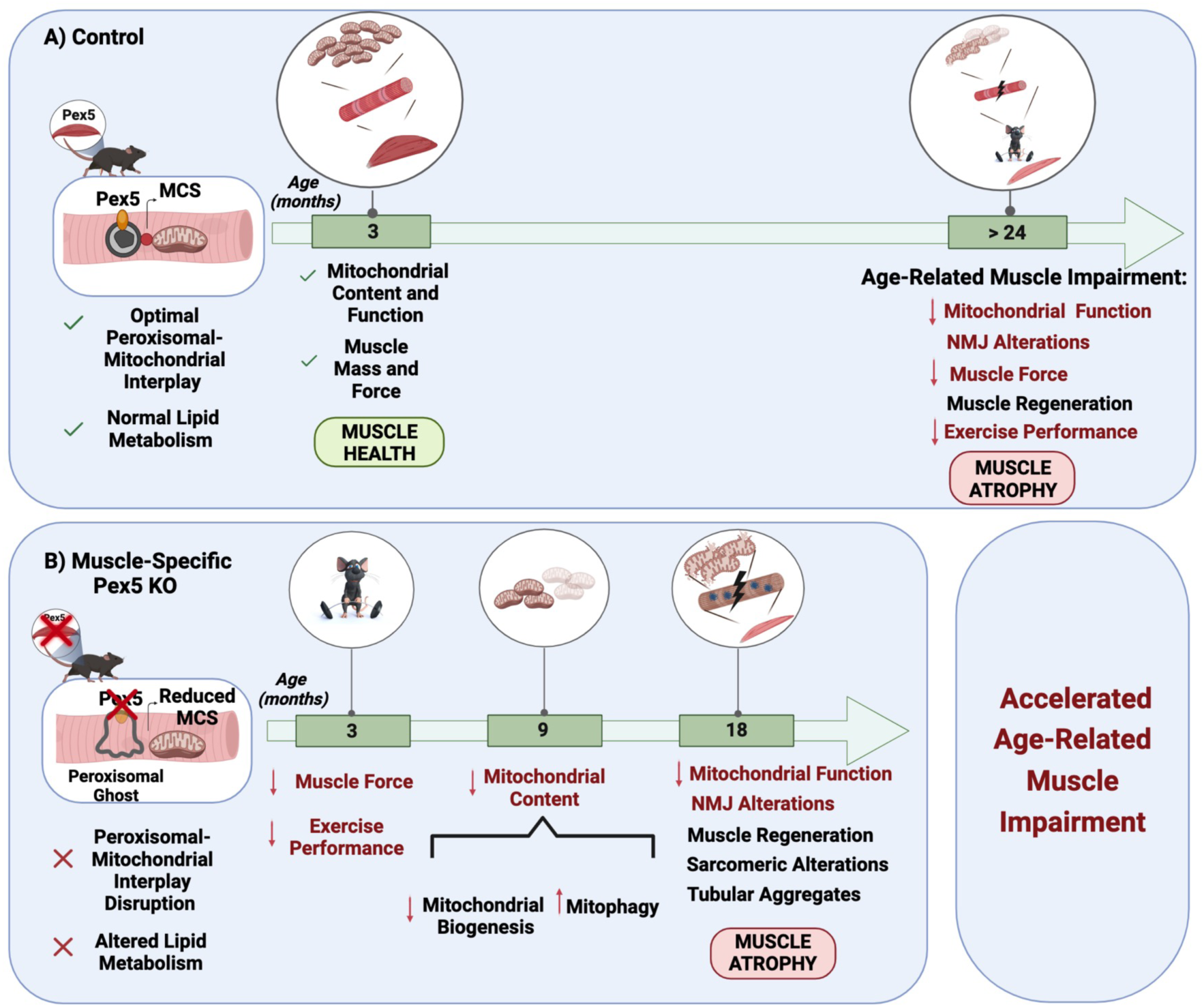
Pex5 is critical to maintain. peroxisomal function and their interplay with mitochondria to preserve muscle force, mass, integrity, and innervation during the aging process. Scheme illustrating the role of Pex5 in skeletal muscle homeostasis. A) In control mice, the age-related defects in muscle structure, metabolism, force, and mass are mainly observed after 24 months of age. B) Pex5 deletion in skeletal muscle leads to abnormal peroxisomal assembly (peroxisomal ghosts), reduced membrane contact sites (MCS) between peroxisomes and mitochondria and altered lipid metabolism. These changes result in early decline in muscle force and exercise performance by 3 months of age. Subsequently, mitochondrial content is affected at 9 months, followed by dysfunction at 18 months. This accelerates age-related muscle atrophy and impairment, characterized by the accumulation of tubular aggregates, proteolytic breakdown of sarcomeres, and degeneration of the neuromuscular junction at 18 months of age. The alterations occurring earlier in KO mice compared to control mice are highlighted in red. The processes Created with BioRender.com.

## DISCUSSION

Peroxisomal dysfunction has been implicated in the pathogenesis of age-related metabolic disorders, including diabetes, obesity, cancer, and aging sarcopenia, all characterized by muscle atrophy and impaired muscle function^19,20^. Here, we show that Pex5 is a novel player in the regulation of skeletal muscle homeostasis, playing a critical role in maintaining muscle health during aging. Using mice with a muscle-specific deletion of Pex5, we demonstrate that the disruption of peroxisomal protein import results into abnormal peroxisome assembly and decreased pexophagy flux within skeletal muscle, confirming the crucial role of Pex5 as a quality control mechanism to eliminate peroxisomes^27–29^. As a result, dysfunctional peroxisomes displaying alterations in lipid metabolism accumulate in the KO muscle. This is characterized by a shift in the fatty acid composition of phospholipid species, such as phosphatidylcholine (PC), phosphatidylethanolamine (PE), and cardiolipins (CL), towards very long-chain unsaturated fatty acids, while simultaneously reducing phospholipid species with shorter fatty acid chains. Furthermore, there is a decrease in ether lipid species such as plasmalogens. Overall, the lipidomic profile of Pex5-null muscles mirrors the observations made in PBD patient skin fibroblasts^32^.

Plasmalogens and phospholipids such as cardiolipin are important components of mitochondrial membranes^49–51^. Previous studies have highlighted their critical role in the maintenance of mitochondrial membrane structure, fluidity, integrity, consequently regulating the organelle biogenesis, morphology, and function^49,52,53^. In fact, both plasmalogens and cardiolipin have a critical function in the mitochondrial respiratory supercomplexes assembly in the inner mitochondrial membrane^31^ contributing to mitochondrial respiration efficiency. Recent findings also highlight the role of peroxisomal-derived plasmalogens in maintaining mitochondrial homeostasis^50^. Peroxisomal dysfunction resulting from adipose-tissue specific Pex16 deletion or the inhibition of plasmalogen synthesis due to glyceronephosphate O-acyltransferase (GNPAT) knockdown leads to a reduction in mitochondrial content and the formation of elongated, dysfunctional mitochondria. Importantly, the introduction of plasmalogens through supplementation rescues these mitochondrial defects, underscoring their potential therapeutic significance^50^. In line with these findings, Pex5 deletion in skeletal muscle triggers a gradual decline in mitochondrial content. This reduction is caused by the inhibition of PGC1α-dependent mitochondrial biogenesis pathways at both 9 and 18 months of age, together with activation of mitophagy at 9 months. The activation of mitophagy likely serves as a beneficial mechanism. It helps to clear organelles affected by alterations in mitochondrial membrane lipid composition, thus maintaining a healthy mitochondrial population in KO muscle tissue at this age. Conversely, by the time KO mice reach 18 months of age, mitochondrial function and mitochondrial lipid metabolism are compromised. Moreover, between 9 and 18 months, there is a progressive and dynamic morphological adaptation in mitochondrial size characterized by elongation, which serves as a tentative mechanism for enhancing metabolic efficiency^54–56^. Accordingly, unbalanced mitochondrial fusion caused by DRP1 deletion specifically in skeletal muscle results in mtDNA loss, mitochondrial dysfunction and mitophagy impairment^57^.

Furthermore, lipid biosynthetic pathways are highly interconnected, requiring the cooperation of different organelles such as peroxisomes, ER and mitochondria^52^. Therefore, lipid alterations impair the communication between peroxisomes and mitochondria, and the physical connections between these organelles at membrane contact sites are critical for efficient metabolic intermediate transfer to achieve key metabolic processes, such as the breakdown of fatty acids through β-oxidation pathways. A deficiency in peroxisome assembly, which leads to the formation of empty peroxisomal ghosts due to Pex5 ablation in skeletal muscle, disrupts the crucial interaction between peroxisomes and mitochondria. Consequently, the membrane contact sites connecting peroxisomes and mitochondria are diminished as early as 3 months of age, coinciding with alterations in lipid and amino acid metabolism in KO muscle. Increased levels of free amino acids such as glutamine, glutamate, asparagine, and aspartate has been linked to increased protein breakdown in conditions characterized by metabolic dysregulation, muscle weakness, and muscle loss^38,39^. Furthermore, the levels of arginine and lysine, whose reduction has been identified as a metabolic signature of unhealthy aging^58^, were decreased in the KO muscle. The changes on lipid and amino acid metabolism parallel the decline in muscle force and exercise performance, occurring prior to the decrease in mitochondrial content and function, as well as muscle atrophy. Notably, in conditions such as aging sarcopenia and cancer cachexia, muscle weakness precedes muscle atrophy^5,59,60^, indicating that age-related muscle dysfunction depends not only on muscle size but also on muscle quality^61^. Therefore, muscle weakness and exercise intolerance arise from impaired peroxisomal function and its consequent disruption in their communication with mitochondria. Conversely, the progression of muscle loss is intricately linked to time-dependent changes in mitochondrial content, morphology, and function. Accordingly, we and others have consistently demonstrated that alterations in mitochondrial content, shape, or function has detrimental consequences for the maintenance of muscle mass and function^11,62^.

These cumulative defects ultimately lead to muscle atrophy and the development of an earlier sarcopenic phenotype by 18 months. This phenotype is characterized by increased center-nucleated fibers, myofiber damage due to tubular aggregate accumulation, Fbxl22-dependent proteolytic breakdown of sarcomeres impacting structural integrity and sarcomeric Z line function, and neuromuscular junction degeneration. The fact that specific inhibition of Pex5 in skeletal muscle can induce muscle denervation without directly impacting the motor neuron was surprising. This finding contrasts with the traditional belief in the context of PBD, where muscle alterations are typically considered secondary to neurological defects. However, our findings, along with other reports ^14,15,63^ , indicate that peroxisomal metabolic activity is required for muscle innervation and function independently of neurological involvement.

NMJ is the critical region where muscle and nerve communicate, influencing each other. In fact, skeletal muscle has a critical role in determining neuron survival, nerve integrity and functional NMJs maintenance. Numerous observations suggest the significant role of retrograde muscle-to-nerve signaling in NMJ maintenance^64^. For example, mitochondrial dysfunction, exclusively in skeletal muscle fibers, correlates with a marked increase in fiber denervation^64–67^. In addition, plasmalogen deficiency in mice leads to NMJ formation alterations and decreased muscle force^68^. Notably, plasmalogens, enriched in healthy muscles^69^, are reduced in skeletal muscle from PBD patients^70^, which are characterized by mitochondrial myopathy, muscle weakness, and muscle atrophy^14,15,71–73^.

The close association between peroxisomal defects and muscle dysfunction is demonstrated also in in a muscle-specific Pex3 knockdown in a Drosophila model where the disruption of peroxisome biogenesis results in the impairment of various processes reliant on muscle function, including eclosion, wing expansion, and climbing^63^.

Further evidence supporting the concept of accelerated age-related skeletal muscle decline in the absence of Pex5 is evident in the lipid profile of 18-month KO muscle, which reflects the alterations observed in sarcopenic muscles. Specifically, the observed increase in total phosphatidylethanolamine (PE) and sphingomyelin (SM) levels in 18-month-old KO muscle aligns with findings in sarcopenic muscle from older subjects and older mice, which exhibit elevated SM levels and significant changes in skeletal muscle phospholipid composition^74–76^. Notably, in this context, PE levels negatively correlate with muscle mass and function^74^. In addition, like 18 months-old KO muscle, sarcopenic muscles display increased levels of long and very long-chain ceramides^77^. The contribution of these lipids to the pathogenesis of age-related muscle disease is highlighted by a study showing that inhibiting ceramide synthesis preserves muscle function during aging^77^, and maintains muscle mass in Colon-26 carcinoma tumor-bearing mice^78^. Moreover, Pex5 transcript and protein levels are reduced in the cortical neurons of aged mice^79^. Similarly, in *C. elegans,* levels of PRX-5 (Pex5 in mice and humans), and nearly 30 other peroxisomal proteins involved in peroxisomal protein import and function are reduced with aging. This decline is associated with age-related peroxisome protein import impairment^80^. Moreover, the peroxisomal protein import is compromised not only in natural but also in an accelerated aging model such as the Hutchinson-Gilford Progeria Syndrome^81,82^. Conversely, dietary fatty acids such as mono-unsaturated fatty acids (MUFA) are linked to longevity by upregulating both peroxisomal and lipid droplet number. Notably, PRX-5 inhibition abolishes MUFA-induced longevity in *C.elegans*^83^.

In summary, our muscle-specific peroxisomal deficient mouse model underscores the importance of preserving peroxisomal function and their interplay with mitochondria to maintain muscle force, integrity, and innervation during the aging process. These findings may constitute the basis for identifying novel mechanisms fundamental to developing drug therapies aimed at preserving muscle function and enhancing the quality of life for individuals affected by peroxisomal disorders and age-related metabolic conditions.

## ACKNOWLEDGEMENTS

We are grateful to Myriam Baes for the *Pex5*-floxed mouse lines,Asushi Miyawaki for the kind gift of mt-mKeima, Camille Bergoglio and to Mikaël Croyal (Nantes Lipidomic Facility Core) for their excellent technical support. This study was supported by AFM Research Grant 24465 and Telethon Research Grant GMR22T1055 to V.R.

## MATERIALS AND METHODS

### Animal handling and generation of muscle-specific Pex5 knockout mice

Animals were handled by specialized personnel under the control of inspectors of the Veterinary Service of the Local Sanitary Service (ASL 16 - Padova), the local officers of the Ministry of Health. All procedures are specified in the projects approved by the Italian Ministero della Salute, Ufficio VI (authorization number 328/2021 PR and 572/2021 PR) and were in compliance with the National Institutes of Health Guidelines for Use and care of Laboratory Animals and with the 1964 Declaration of Helsinki and its later amendments. To generate constitutive muscle-specific Pex5 knockout animals, mice bearing Pex5 floxed alleles^84^ (Pex5f/f) (provided by Myriam Baes) were crossed with transgenic mice expressing Cre under the control of a Myosin Light Chain 1 fast promoter (MLC1f-Cre)^85^. Mice were maintained on a 12-hour light/12-hour dark schedule and were fed standard rodent food chow and water ad libitum. Denervation was performed by cutting the sciatic nerve of the left limb, while the right limb was used as control. Surgical procedures, including sciatic nerve cutting and muscle electroporation, were performed under inhalation of isoflurane in medical oxygen with post-operative analgesia administerd using carprofen or meloxicam. Mice were sacrificed by cervical dislocation, and the different tissues were weighed and then frozen in liquid nitrogen, and utilized for histological experiments, immunohistochemistry or gene expression studies. Experiments were performed on 3, 9 and 18 months-old adult male mice. Cre-negative littermates were used as controls.

### Body composition analyses

Quantitative magnetic resonance was utilized to measure lean and fat mass in live mice using the EchoMRITM-100 system (EchoMRI LLC).

### Exercise studies

Mice aged 3 and 18 months-old were acclimated to and trained on a treadmill with no inclination (Biological Instruments, LE 8710 Panlab Technology 2B) over a period of three days. During these 3 consecutive days preceding the test, mice ran for 5 min at 10 m/min. On the fourth day, mice underwent a single bout of running starting a speed of 10 m/min. Forty minutes later, the treadmill speed was increased by 1 m/min every 10 min for a total of 30 min, followed by an increase of 1 m/min every 5 min until mice were exhausted. Exhaustion was defined as the point at which mice spent more than 5 seconds on the electric shocker without attempting to resume running. Total running total running distance was calculated for each mouse.

### Gene expression analyses

Total RNA was prepared from tibialis anterior muscles using TRIzol (Life Technologies). Complementary DNA was generated from 0.4 μg of RNA reverse-transcribed with SuperScript III Reverse Transcriptase (Invitrogen). Duplicates of cDNA samples were then amplified on the 7900HT Fast Real-Time PCR System (Applied Biosystems, MA, U.S.) using the Power SYBR Green RT-PCR kit (Applied Biosystems). The relative expression ratio of target gene was calculated based on PCR efficiency and quantification cycle deviation (ΔCq) of an unknown sample versus a control, and expressed in comparison to the reference gene^86^. All data were normalized to GAPDH or HPRT expression, of which abundance did not change under any of the experimental conditions, and plotted in arbitrary units as mean ± SEM. The sequences of the oligonucleotide primers used are listed in Table S1.

### RNA sequencing and analysis

Total RNA was isolated from gastrocnemius muscle using TRIzol reagent (Life Technologies) according to the manufacturer’s instructions. The isolated RNA was initially assessed for purity using a Qubit fluorometer (Thermo Fisher Scientific) and it was then submitted to the Biology Department at the University of Padua. The RNA was assessed for integrity using the TapeStation 4150 (Agilent). Subsequently, the QuantSeq 3’ mRNA-seq Library Prep kit (Lexogen) was employed for library construction. The library generation process is initiated with oligo(dT) priming, where the primer already contained Illumina-compatible linker sequences. After first strand synthesis the RNA was removed, and second strand synthesis was initiated by random priming and a DNA polymerase. The random primer also contains Illumina-compatible linker sequences. Second strand synthesis was followed by a magnetic bead-based purification step. The library was then amplified, introducing the sequences required for cluster generation. External barcodes were introduced during the PCR amplification step. Library quantification was performed by fluorometer (Qubit) and TapeStation 4150 (Agilent). QuantSeq Forward contains the Read 1 linker sequence in the second strand synthesis primer, hence NGS reads were generated towards the poly(A) tail and directly corresponded to the mRNA sequence. QuantSeq FWD maintains strand-specificity and allows mapping of reads to their corresponding strand on the genome, enabling the discovery and quantification of antisense transcripts and overlapping genes. Sequencing was performed on NextSeq500 ILLUMINA instrument to produce at least 10 million of reads (75bp SE) for sample.

### Immunohistochemistry and stainings

Cryosections of adult tibialis anterior muscle were stained for Hematoxylin & Eosin (H&E), Succinate dehydrogenase (SDH), and modified Gomori trichrome staining. For fiber typing, TA slides were incubated with the following monoclonal antibody combination: BA-D5 (1:100) (Developmental Studies Hybridoma Bank) for the type 1 MyHC isoform, SC-71 (1:100) (Developmental Studies Hybridoma Bank) for the type 2A MyHC isoform and anti-Dystrophin (1:100) (ab15277, Abcam. Cambridge, VK) for the sarcolemma. Fibers negative for BA-D5 and SC-71 were identified as Type 2X and 2B. Images were captured using a Leica DFC300-FX digital charge-coupled device camera and the Leica DC Viewer software. Cross-sectional area (CSA) was calculated measuring the cross-sectional area of all individual fibers from entire muscle cross-section from tibialis anterior muscles based on assembled mosaic image (20× magnification). The morphometric analyses were made using MATLAB Semi-Automatic Muscle Analysis using Segmentation of Histology (SMASH) software. The specific antibodies used for immunostaining are listed in Table S2.

### NMJ measurements

EDL muscles from 18-month-old control and KO mice were dissected and fixed in 4% PFA in PBS for 30 min at room temperature. Subsequently, muscle fiber bundles were teased by mechanical manipulation in PBS with the aid of a stereomicroscope and then quenched in 0.24% NH_4_Cl PBS for 20 minutes. Following permeabilization and a 2-hour saturation period in blocking solution (15% goat serum, 2% BSA, 0.25% gelatine, 0.20% glycine, 0.5% Triton-X100 in PBS), samples were incubated with a primary antibody against VAMP1 (1:200) in blocking solution at 4 °C for 72 h. Muscle fibers were then washed and incubated with secondary antibodies and α-BTx AlexaFluor-555 (1:200) (ThermoFisher B35451) to label post-synaptic acetylcholine receptors (AChRs). Images were acquired using a Zeiss LSM 900 Confocal microscope equipped with a 40× HCX PL APO NA 1.4 oil immersion objective. To minimize crosstalk, laser excitation line, power intensity, and emission range were optimized for eachs specific fluorophore of interest. Recovery extent was assessed by determining the proportion of denervated, partially denervated, and innervated NMJs through analysis of fluorescence signal overlap between pre-synaptic and post-synaptic staining, employing Fiji’s plugin called ColocThreshold. Arbitrary thresholds were set to evaluate NMJ innervation status: NMJs with an overlap range of 0-40%, indicating absent or reduced pre-synaptic staining, were classified as completely denervated, those with a range of 40-80% were considered partially denervated, while those above 80% were classified as innervated.

### Transmission Electron Microscopy (EM)

For EM, Extensor Digitorum Longus (EDL) muscles were dissected from sacrificed animals, pinned on a Sylgard dish, fixed at room temperature with 3.5% glutaraldehyde in a 0.1 M NaCaCO buffer (pH 7.4), and stored in the fixative solution at 4 °C. Fixed muscles were then post-fixed in a mixture of 2% OsO4 and 0.8% K3Fe(CN)6 for 1–2 h, rinsed with a 0.1 M sodium cacodylate buffer with 75 mM CaCl2, en-bloc stained with saturated uranyl acetate, and embedded for EM in epoxy resin (Epon 812) as in^87^. Ultrathin sections (∼40 nm) were cut in a Leica Ultracut R microtome (Leica Microsystem, Austria) using a Diatome diamond knife (Diatome Ltd, CH-2501 Biel, Switzerland) and examined at 60 kV after double-staining with uranyl acetate and lead citrate, with an FP 505 Morgagni Series 268D electron microscope (FEI Company, Brno, Czech Republic), equipped with Megaview III digital camera (Munster, Germany) and Soft Imaging System (Germany). Mitochondrial area (Fig.4B) was measured in the same set of micrographs using Image J. Only mitochondria which were entirely visualized in the micrograph were measured.

### Measurement of peroxisome number, morphology, and size

Confocal images of isolated FDB muscle fibers were obtained using a Zeiss LSM900 upright confocal microscope equipped with a Plan-Neofluar 40×/1.3 oil-immersion objective (Carl Zeiss). To analyze the number, morphology, and size of PMP70-positive structures in both control and KO FDB isolated fibers, we used the Analyze Particles function (ImageJ). The resulting parameters results were normalized to muscle fiber area.

### Force measurements

In vivo gastrocnemius force measurements were performed as described previously^88^. Briefly, mice were anesthetized, and stainless-steel electrodes wires were placed on either side of the sciatic nerve. Torque production of the plantar flexors was measured using a muscle lever system (Model 305c; Aurora Scientific, Aurora ON, Canada). The force– frequency curves were determined by increasing the stimulation frequency in a stepwise manner, pausing for 30s between stimuli to avoid effects due to fatigue. Following force measurements, animals were sacrificed by cervical dislocation and muscles were dissected and weighted. Force was normalized to the gastrocnemius muscle mass as an estimate of specific force.

### Mitochondrial isolation

Mitochondria from quadriceps muscles of the indicated genotypes were isolated as described and were then probed with the indicated antibodies by immunoblotting.

### Measurement of mitochondrial DNA copy number

The total tibialis anterior DNA was isolated using Puregene Cell and Tissue Kit (Qiagen). The content of mtDNA was calculated using real-time quantitative PCR (qRT-PCR) by measuring the threshold cycle ratio (ΔCt) of the mitochondrial-encoded gene CoxI versus the nuclear-encoded gene RNaseP. The following primers were used:

Cox1_fw: 5′-TGCTAGCCCGACAGGCATTACT-3′

Cox1_rv: 5′-CTGACCACACGAGCTGGTAGAA-3′

RNaseP_fw: 5′-GCCTACACTGGAGTCGTGCTACT-3′

RNaseP_rv: 5′-CGGGATCAAAGAAAGTTGTGTTT-3′

### OCR (oxygen consumption rate) measurements

Single isolated FDB fibers, plated on laminin-coated XF24 microplate wells and cultured in DMEM (D5030 Sigma-Aldrich), supplemented with 1 mM NaPyr, 5 mM glucose, 33 mM NaCl, 15 mg phenol red, 25 mM HEPES, and 1 mM of L-Glu. Fibers were maintained in culture for 2 h at 37 °C in 5% CO2. The rate of oxygen consumption was assessed in real-time with the XF24 Extracellular Flux Analyzer (Agilent), which allows to measure OCR changes following up to four sequential additions of compounds. A titration with the uncoupler CCCP was performed to determine the optimal CCCP concentration (0.6 μM) that maximally increases OCR. To calculate basal and maximal respiration, non-mitochondrial O2 consumption was substracted from absolute values. ATP-linked respiration was calculated as the difference between basal and oligomycin-insensitive O2 consumption. The results were normalized to the fluorescence of Calcein (Sigma-Aldrich). Fibers were loaded with 2 μM Calcein for 30 min. Fluorescence was measured using a Perkin Elmer EnVision plate reader in well scan mode using a 480/20 nm filter for excitation and 535/20 nm filter for emission.

### Mitochondrial respiratory chain enzymatic activity

Assessment of mitochondrial respiratory chain enzymatic activities on muscles was determined as described previously^89^. Results were normalized to citrate synthase enzymatic activity (Fig.3G and 4D) or to total protein content (Suppl. Fig.4B).

### Immunoblotting

To obtain whole skeletal muscle lysates, samples were homogenized in lysis buffer containing 50 mM Tris (pH 7.5), 150 mM NaCl, 5 mM MgCl2, 10% glycerol, 1% SDS, 1% Triton X-100, and inhibitors for phosphatase (P5726, Sigma) and protease (P8340, Sigma). Protein fractions were resolved by SDS-PAGE using pre-cast 4-12% Bis-Tris gels (Thermo Fisher Scientific), blotted onto nitrocellulose membranes (BioRad), and incubated with the appropriate primary antibody overnight. Blots were stripped 2 × 10 min washes in mild stripping buffer containing 0.2 M Glycine, 0.1% SDS, 1% Tween-20 (pH 2.2) and reprobed if necessary. The membranes were visualized with the ImageQuant LAS 4000 and quantified using ImageJ software (https://imagej.nih.gov/ij/). Protein expression was normalized to GAPDH. In Fig. S3B, protein expression of isolated mitochondria was normalized to total protein content revealed with Ponceau S staining. List of antibodies is provided in Table S2.

### Autophagic flux quantification

We monitored autophagic flux in basal conditions using colchicine (Sigma-Aldrich Chemie, C9754)^90^ . Briefly, 3 months-old mice were treated with 0,4mg/kg of colchicine or vehicle by intraperitoneal injection. The treatment was administered twice, at 24 h and at 12 h before muscle dissection. Total muscle homogenates were used to measured general autophagy and pexophagy flux respectively.

### *In Vivo* FDB Electroporation

Electroporation experiments were performed on FDB muscles from control and knockout animals. The animals were anesthetized by inhalation of isoflurane in medical oxygen. Ten microliters of Hyaluronidase (2 mg/mL) (Sigma-Aldrich, St. Louis, MO, USA) were injected in the feet of anesthetized mice to soften muscle tissue underneath the epidermis. After 50 min, we injected 10 μg of plasmid DNA (mitochondria-targeted mKeima (mt-mKeima) or PO/MT SPLICS-P2A), and after 10 min, electric pulses were applied by two stainless needles placed at 1 cm from each other (100 Volts/cm, 20 pulses, 1 s intervals). Muscles were analyzed 10 days later. FDB muscles were collected in 1% P/S DMEM. No evidence of necrosis or inflammation was observed after the transfection procedure. FDBs were digested in type I collagenase at 37 °C for 1,5-2 hours. The fibers were dissociated by creating mechanical forces using a pipette. The single isolated fibers were then plated on glass coverslips coated with 10% Matrigel in Tyrode’s salt solution (pH 7.4) and were used to monitor mitophagy and peroxisome-mitochondria contact sites in transfected FDB single fibers.

### Imaging and quantification of peroxisome-mitochondria contact sites

PO/MT SPLICS-P2A probe^36^ was used to investigate peroxisome-mitochondria contact sites in transfected FDB single fibers. Muscle fibers were imaged with a using a Zeiss LSM900 upright confocal using a Plan-Neofluar 40×/1.3 oil-immersion objective (Carl Zeiss). Images were acquired by using the Zeiss software. The SPLICS signal is acquired at lasers wavelength of 488nm. Z-stack of the muscle sections were acquired every 0,5 μm, then processed using ImageJ (National Institutes of Health (NIH)). Images were first convolved and selected by freehand selection of ImageJ in the drawing/selection polygon tool and then processed using the “Quantification 1” plugin (https://github.com/titocali1/Quantification-Plugins). A 3D reconstruction of the resulting image was obtained using the VolumeJ plugin (https://github.com/titocali1/Quantification-Plugins). A selected face of the 3D rendering was then thresholded and used to count short contact sites through the “Quantification 2” plugin (https://github.com/titocali1/Quantification-Plugins). Contacts are measured as number of contacts normalized to fiber area (contacts/µm^2^).

### Mito-mKeima mitophagy assay

Mt-mKeima was used to monitor mitophagy in transfected FDB single fibers. Mt-mKeima is a coral-derived protein that exhibits both pH-dependent excitation and resistance to lysosomal proteases. These properties allow for rapid determinations as to whether the protein is in mitochondria or in the lysosome. In fluorescence microscopy, ionized Keima is detected as a red fluorescent signal at lower pH values (lysosome) and neutral Keima is detected as a green fluorescent signal at higher pH values (mitochondria). Fluorescence of mt-mKeima was imaged in two channels via two sequential excitations (458 nm, green; 561 nm, red) and using a 570-to-695 nm emission range^91,92^. The level of mitophagy was defined as the total number of red pixels divided by the total number of all pixels.

### Metabolomics

Metabolomics was performed as previously described, with minor adjustments^93^. In a 2 mL tube, the following amounts of internal standard dissolved in water were added to each sample of freeze-dried gastrocnemius muscle: adenosine-^15^N_5_-monophosphate (5 nmol), adenosine-^15^N_5_-triphosphate (5 nmol), D_4_-alanine (0.5 nmol), D_7_-arginine (0.5 nmol), D_3_-aspartic acid (0.5 nmol), D_3_-carnitine (0.5 nmol), D_4_-citric acid (0.5 nmol), ^13^C_1_-citrulline (0.5 nmol), ^13^C_6_-fructose-1,6-diphosphate (1 nmol), ^13^C_2_-glycine (5 nmol), guanosine-^15^N_5_-monophosphate (5 nmol), guanosine-^15^N_5_-triphosphate (5 nmol), ^13^C_6_-glucose (10 nmol), ^13^C_6_-glucose-6-phosphate (1 nmol), D_3_-glutamic acid (0.5 nmol), D_5_-glutamine (0.5 nmol), D_5_-glutathione (1 nmol), ^13^C_6_-isoleucine (0.5 nmol), D_3_-lactic acid (1 nmol), D_3_-leucine (0.5 nmol), D_4_-lysine (0.5 nmol), D_3_-methionine (0.5 nmol), D_6_-ornithine (0.5 nmol), D_5_-phenylalanine (0.5 nmol), D_7_-proline (0.5 nmol), ^13^C_3_-pyruvate (0.5 nmol), D_3_-serine (0.5 nmol), D_6_-succinic acid (0.5 nmol), D_4_-thymine (1 nmol), D_5_-tryptophan (0.5 nmol), D_4_-tyrosine (0.5 nmol), D_8_-valine (0.5 nmol). Subsequently, solvents were added to achieve a total volume of 500 µL methanol and 500 µL water. A 5 mm stainless steel bead was added and a Qiagen TissueLyser II was used for 5 minutes at 30 times/s to homogenize each sample, before the addition of 1 mL chloroform. After thorough mixing, samples were centrifuged for 10 min at 14.000 rpm. The polar top layer was transferred to a new 1.5 mL tube and dried using a vacuum concentrator at 60°C. Dried samples were reconstituted in 100 µL 6:4 (v/v) methanol:water. Metabolites were analyzed using a Waters Acquity ultra-high performance liquid chromatography system coupled to a Bruker Impact II™ Ultra-High Resolution Qq-Time-Of-Flight mass spectrometer. Samples were kept at 12°C during analysis and 5 µL of each sample was injected. Chromatographic separation was achieved using a Merck Millipore SeQuant ZIC-cHILIC column (PEEK 100 x 2.1 mm, 3 µm particle size). Column temperature was held at 30°C. Mobile phase consisted of (A) 1:9 (v/v) acetonitrile:water and (B) 9:1 (v/v) acetonitrile:water, both containing 5 mmol/L ammonium acetate. Using a flow rate of 0.25 mL/min, the LC gradient consisted of: Dwell at 100% Solvent B, 0-2 min; Ramp to 54% Solvent B at 13.5 min; Ramp to 0% Solvent B at 13.51 min; Dwell at 0% Solvent B, 13.51-19 min; Ramp to 100% B at 19.01 min; Dwell at 100% Solvent B, 19.01-19.5 min. Column was equilibrated by increasing flow rate to 0.4 mL/min at 100% B for 19.5-21 min. MS data were acquired using negative and positive ionization in full scan mode over the range of m/z 50-1200. Data were analyzed using Bruker TASQ software version 2021.1.2.452. All reported metabolite intensities were normalized to the sum of all adenosine nucleotides, as well as to internal standards with comparable retention times and response in the MS. Metabolite identification has been based on a combination of accurate mass, (relative) retention times, ion mobility data and fragmentation spectra, compared to the analysis of a library of standards.

### Lipidomics

Lipidomics analysis was performed as previously described^32^. In a 2 ml tube freeze-dried gastrocnemius muscle was added and mixed with a mix of internal standards for different lipid classes, including 0.1 nmol of cardiolipin CL(14:0/14:0/14:0/14:0), 2.0 nmol of phosphatidylcholine PC(14:0/14:0), 0.1 nmol of phosphatidylglycerol PG(14:0/14:0), 5.0 nmol of phosphatidylserine PS(14:0/14:0), 0.5 nmol of phosphatidylethanolamine PE(14:0/14:0), 0.5 nmol of phosphatidic acid PA(14:0/14:0), 2.125 nmol of sphingomyelin SM(d18:1/12:0), 0.02 nmol of lysophosphatidylglycerol LPG(14:0), 0.1 nmol of lysophosphatidylethanolamine LPE(14:0), 0.5 nmol of lysophosphatidylchloline LPC(14:0), 0.1 nmol of lysophosphatidic acid LPA(14:0), 0.5 nmol of phosphatidylinositol PI(8:0/8:0), 0.5 nmol diglycerides DG(14:0/14:0), 0.5 nmol triglycerides TG(14:0/14:0/14:0), 2.5 nmol cholesterol ester D7-CE(16:0), 0.125 nmol of sphingosine and ceramide mix (Avanti Polar Lipids) dissolved in 1:1 (v/v) methanol:chloroform. Next, 1.5 ml 1:1 (v/v) methanol:chloroform was added to each sample. The mixture was sonicated in a water bath (5 min) and centrifuged (4 °C, (16,000×g, 10 min). The supernatant was transferred to a 1.5 ml glass auto sampler vial and evaporated under a stream of nitrogen at 45°C. The dried lipids were reconstituted in 100 μl of 1:1 (v/v) chloroform:methanol. Chromatographic separation of lipids was done using a Thermo Fisher Scientific Ultimate 3000 binary UPLC using a normal phase and a reverse phase column in separate runs. Normal-phase separation was done using a Phenomenex® LUNA silica, 250×2 mm, 5 µm 100 Å column. Column temperature was held constant at 25°C. The composition of the mobile phase A consisted of 85:15 (v/v) methanol:water containing 0.0125% formic acid and 3.35 mmol/l ammonia and the composition of mobile phase B consisted of 97:3 (v/v) chloroform:methanol containing 0.0125% formic acid. The LC gradient started at of 10% A for 0-1 min, 20% A at 4 min, 85% A at 12 min, 100% A at 12.1 min, 100% A for 12.1-14 min, 10% A at 14.1 min, 10% A for 14.1-15 min using a flow rate of 0.3 ml/min. Reversed-phase separation was done using a Waters HSS T3 column (150×2.1 mm, 1.8 μm particle size). The composition of the mobile phase A consisted of 4:6 (v/v) methanol:water and B 1:9 (v/v) methanol:isopropanol, both containing 0.1% formic acid and 10 mmol/l ammonia. The gradient started at 100% A going to 80% A at 1 min and 0% A at 16 min, 0% A for 16-20 min, 100% A at 20.1 min and 100% A for 20.1-21 min. The column temperature was held constant at 60°C and a flow rate of 0.4 ml/min was used. After LC separation, lipids were detected using a Q Exactive Plus Orbitrap mass spectrometer (Thermo Scientific) using negative and positive ionization. The spray voltage was 2,500 V and nitrogen was used as the nebulizing gas. A resolution of 280.000 was used in a mass range of m/z 150 to m/z 2.000.

### Targeted Lipidomics

Around 30 mg of muscle tissue from 9 months-old mice for each condition was diluted in 300 µL PBS 1X and tissue homogenized with precellys®. Lipids of supernatant were extracted using the method of Folch-Lees {Folch, 1957 #59}. The extracts were filtered, and lipids recovered in the methanol-chloroform phase. TAG and DAG were isolated using thin layer chromatography on silica glass plates (E. Merck, Darmstadt, Germany) developed in petroleum ether, ethyl ether, acetic acid (80:20:1) and visualized by fluorescein (2,7 – dichlorofluoresceine (Fluka) 0.2% in ethanol). The TAG and DAG bands were scraped from the plate and transmethylated using 5% acetyl chloride/95 % methanol. The methylated fatty acids were extracted with isooctane and analyzed by gas chromatography using a gas chromatograph GC30 equipped with flame ionization detectors (Shimatzu), and CP-Wax 58 capillary column, 50 m in length,0.25 mm external diameter, 0.2 μm thickness of the stationary phase (Varian Inc., Les Ulis, France)). Helium was used as a carrier gas. The oven temperature was: initial temperature of 60 °C – 110 °C at a rise rate of 20 °C · min^−1^, then 223 °C at a rate of 1.6 °C · min^−1^, 2 minutes at this temperature, then 230 °C at a rate of 2 °C · min^−1^, 13 minutes at this temperature, then 270 °C at a rate of 40 °C · min^−1^, then remaining at this temperature.. Fatty acid methyl esters are identified by comparing the retention times to those of known standards. Inclusion of the internal standard, trinonadecanoyl glycerol and diheptadecanoyl glycerol (Sigma), permits quantitation of the amount of TAG and DAG in the sample.

### Statistical analysis

Statistical tests (Welch t test, Student’s t test or Mann-Whitney test, or two-way ANOVA, and the multiple testing procedures) were used as described in the figure legends and were conducted upon verification of the normality assumption using the Shapiro-Wilk test (where applicable). For all graphs, data are presented as means ± SEM. Differences between groups were considered statistically significant when the P value obtained was less than 0.05; the P values are reported in the figure legends. Statistical analyses were performed using GraphPad Prism 7.0a (GraphPad).

