## Supplementary Figures and Tables for "ALTERATIONS IN PEROXISOMAL-MITOCHONDRIAL INTERPLAY IN SKELETAL MUSCLE ACCELERATES MUSCLE DYSFUNCTION"

**Supplementary Figure 1**

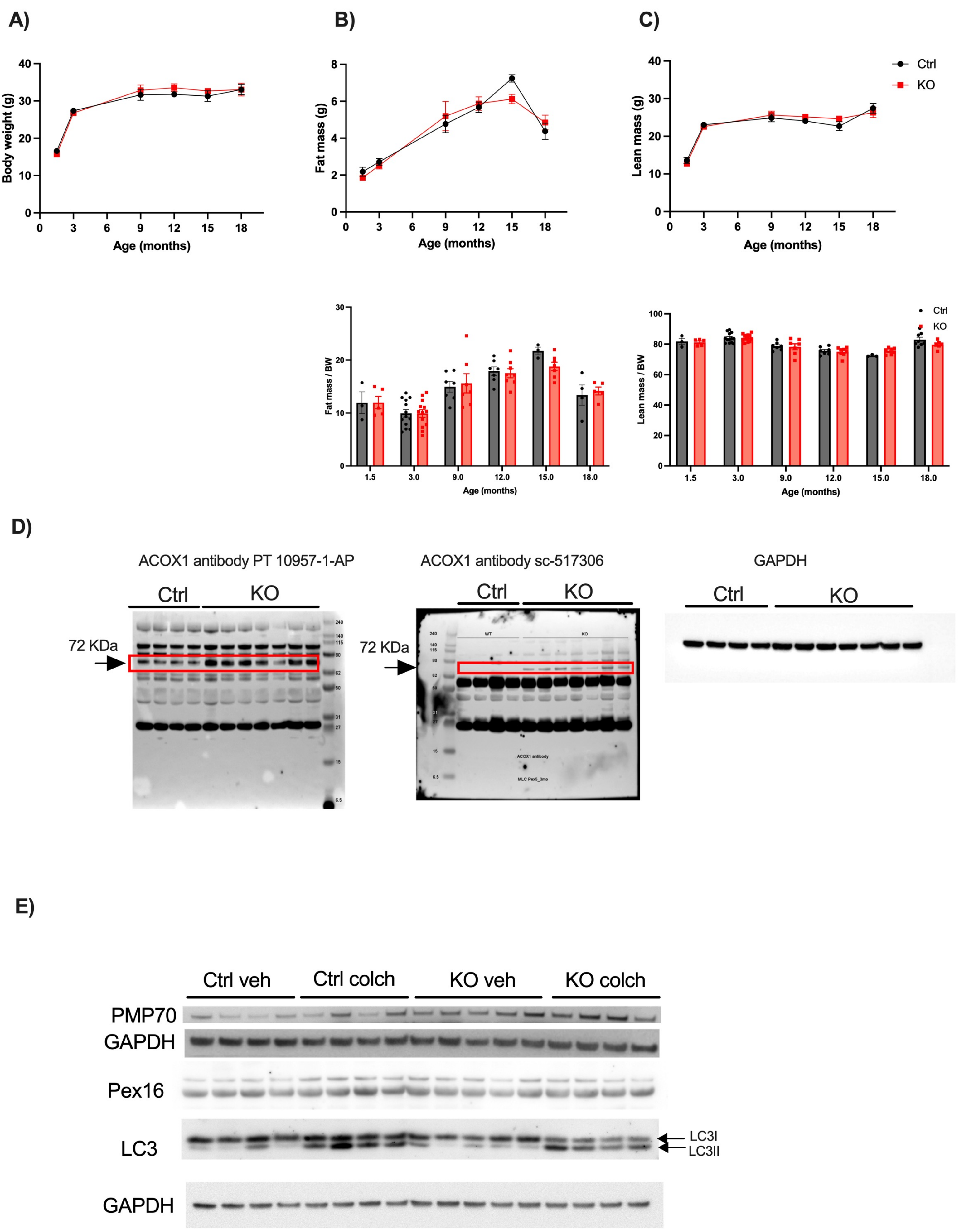

#### Supplementary Figure 2

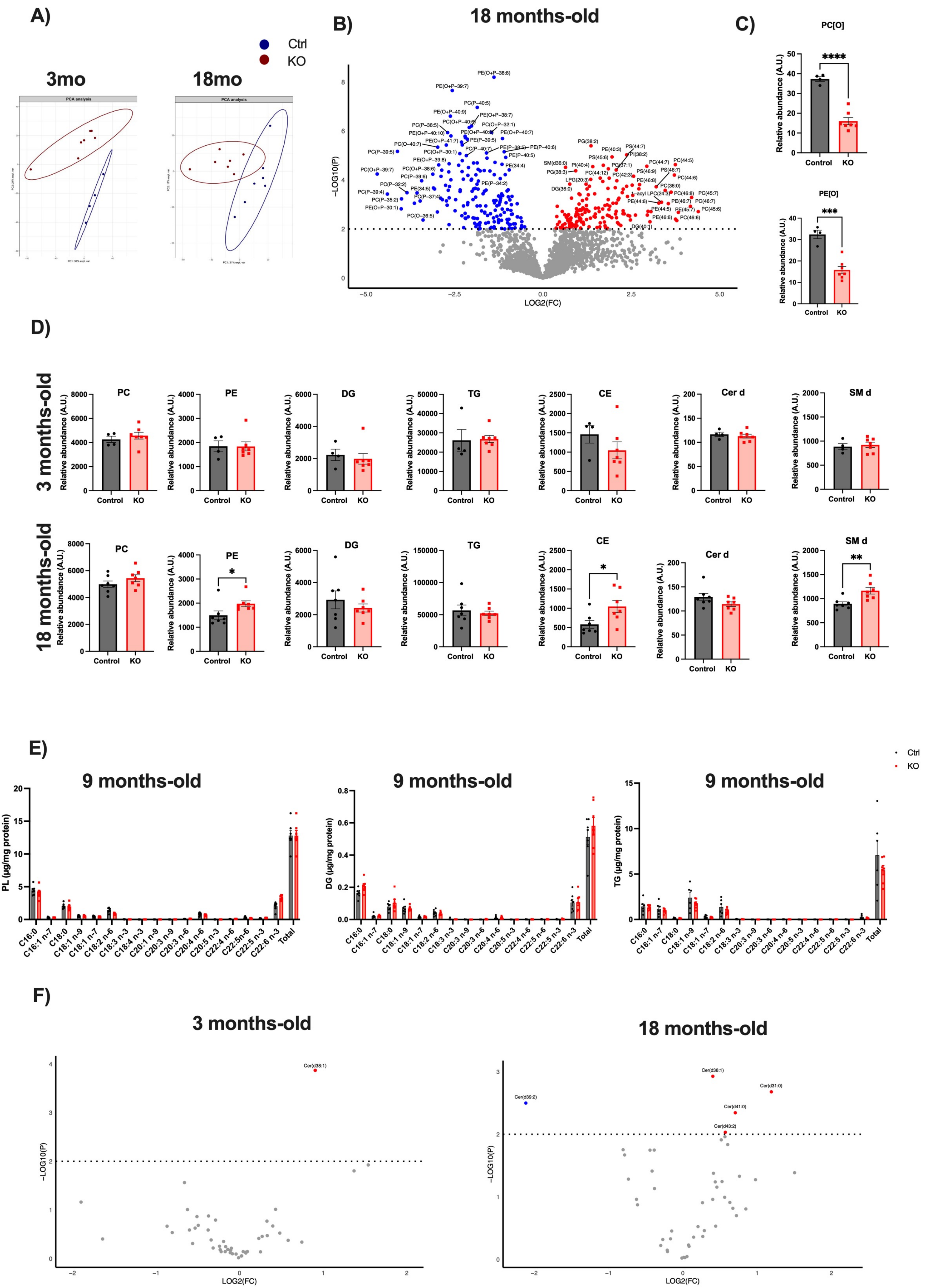

#### Supplementary Figure 3

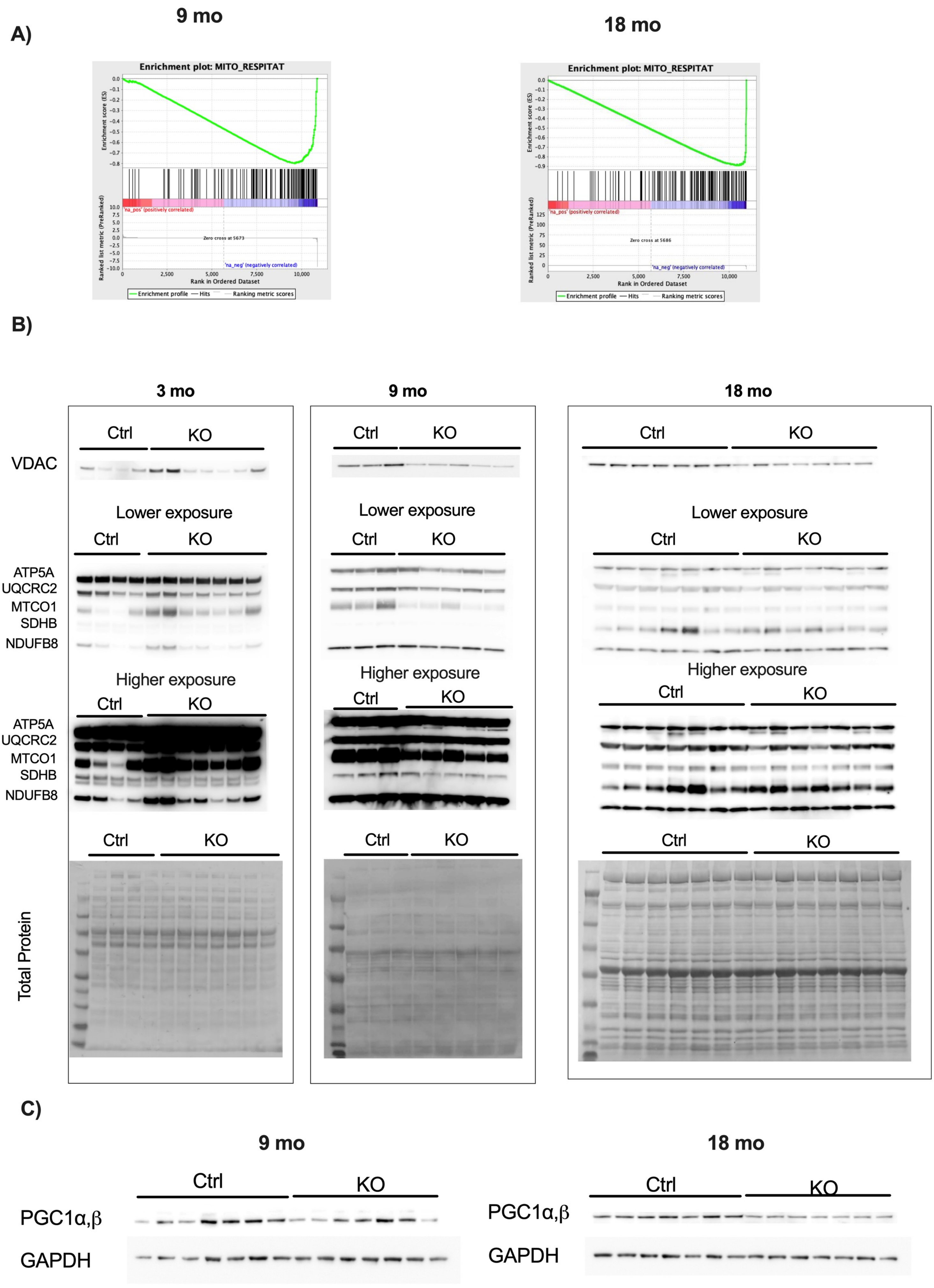

#### Supplementary Figure 4

A)

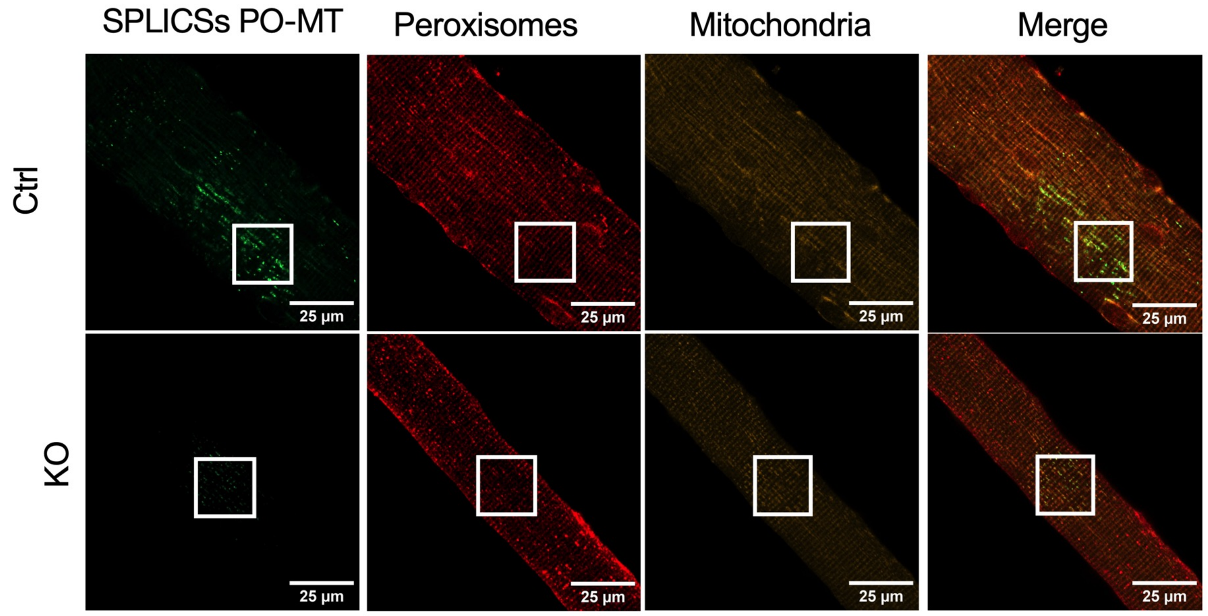


B)

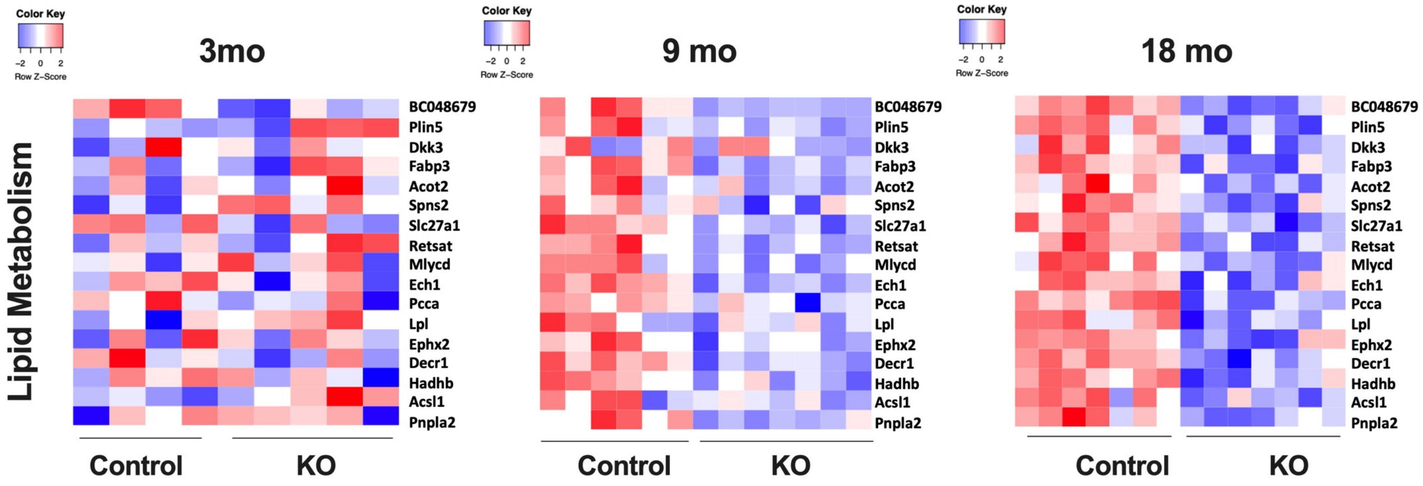


C)

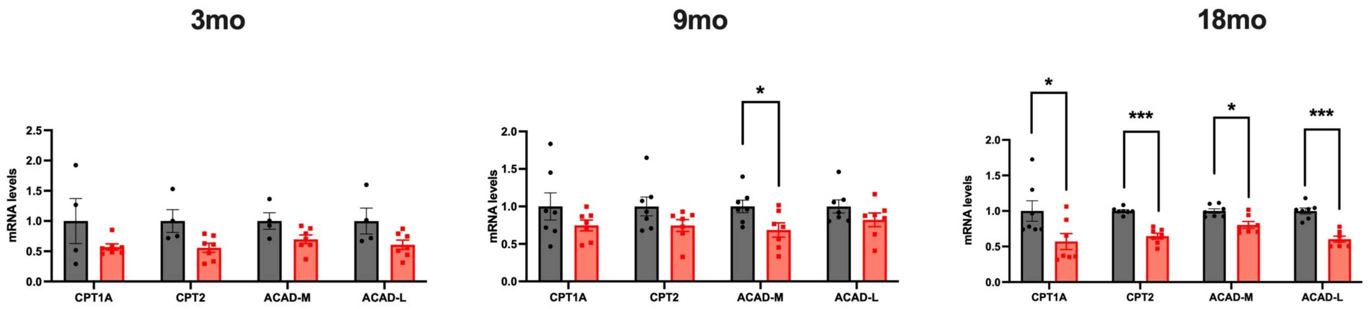

### Supplementary Figure 5

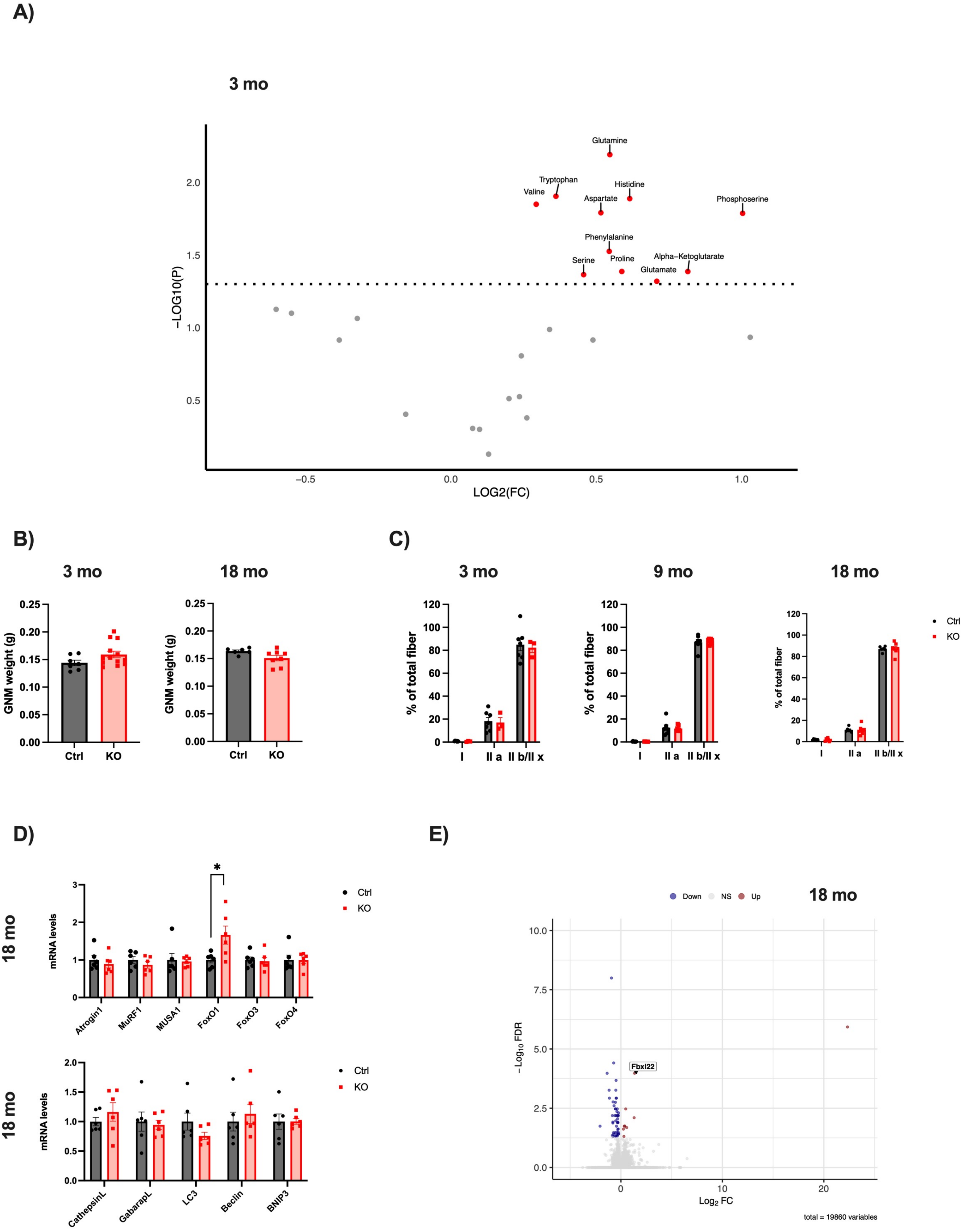

#### Supplementary Figure 6

A)

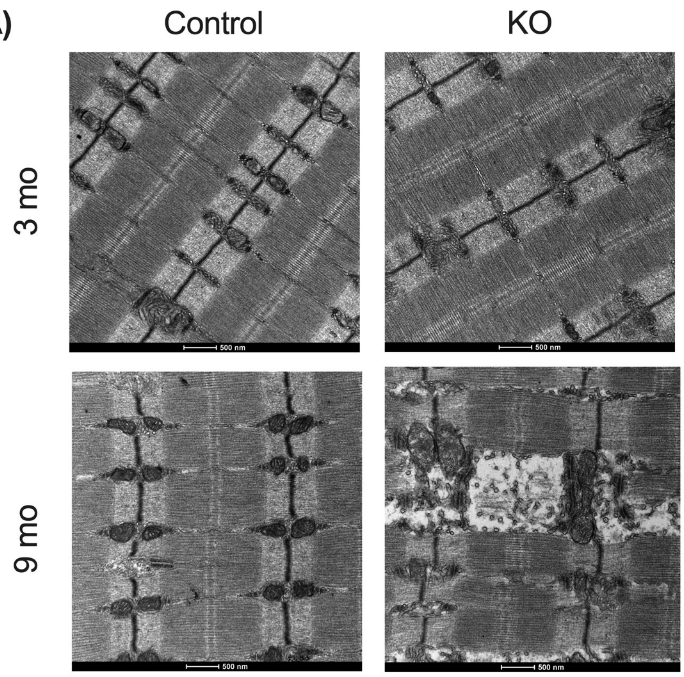


B)

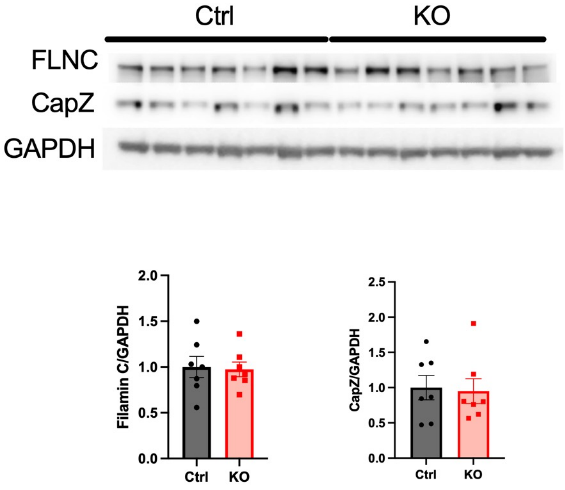


C)

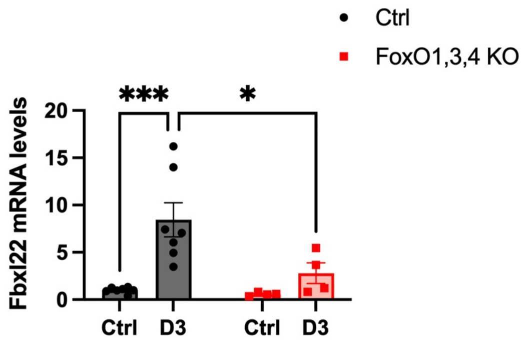


D)

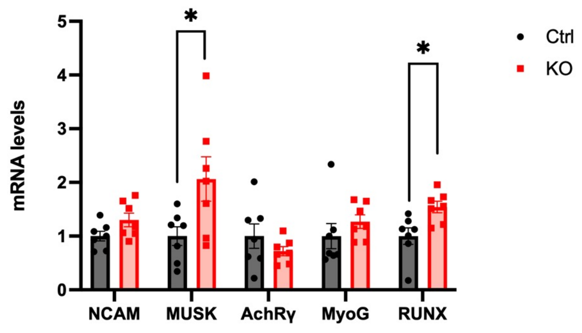

**Table S1.** Real time primer sequences

| *Gene name* | *Forward primer* | *Reverse primer* |
|---|---|---|
| *GAPDH* | CACCATCTTCCAGGAGCGAG | CCTTCTCCATGGTGGTGAAGAC |
| *PEX5* | TCAAACTTTCGCCAGGCTCC | GAATTCTTGGGACCAGTCGGT |
| *ATROGIN1* | GCAAACACTGCCACATTCTCTC | CTTGAGGGGAAAGTGAGACG |
| *MURF1* | ACCTGCTGGTGGAAAACATC | CTTCGTGTTCCTTGCACATC |
| *MUSA1* | TCGTGGAATGGTAATCTTGC | TCGTGGAATGGTAATCTTGC |
| *FOXO1* | GCTGGGTGTCAGGCTAAGAG | AGGGGTGAAGGGCATCT |
| *FOXO3* | CTGTGGCTGAGTGAGTCTGAA | ACCTTCGTCTCTGAACTCCTT |
| *FOXO4* | CGGAGTGAAAGGGACAGTTTAG | CCCTGTGGCTGACTTCTTATTC |
| *CATHEPSIN-L* | GTGGACTGTTCTCACGCTCAAG | TCCGTCCTTCGCTTCATAGG |
| *GABARAP-L* | CATCGTGGAGAAGGCTCCTA | ATACAGCTGGCCCATGGTAG |
| *LC3* | CACTGCTCTGTCTTGTGTAGGTTG | TCGTTGTGCCTTTATTAGTGCATC |
| *BECLIN1* | TGGAAGGGTCTAAGACGT | GGCTGTGGTAAGTAATGGA |
| *BNIP3* | TTCCACTAGCACCTTCTGATGA | GAACACCGCATTTACAGAACAA |
| *COX-I* | TGCTAGCCCGACAGGCATTACT | CTGACCACACGAGCTGGTAGAA |
| *RNAseP* | GCCTACACTGGAGTCGTGCTAGT | CGGGATCAAAGAAAGTTGTGTTT |
| *CPT1A* | GAAGAAGAAGTTCATCCGATTCAAG | CGCCACTCACGATGTTCTT |
| *CPT2* | GCCAGTTCAGGAAGACAGA | GCAGAAACAGTCAAGTTGGTG |
| *ACAD-M* | CCAATGATGTGTGCTTACTGTG | GGGTACTTTAGGATCTGGGTTA |
| *ACAD-L* | GGCTGGTTAAGTGATCTCGTG | CTGGCAATCGGACATCTTC |
| *FBXL22* | TGGCATTCCAGCCGTGT | GTGACGGACGTCAGGTT |
| *NCAM* | ACAATGCTGCGAACTAAGGA | TGCCACTTCACACACAGGA |
| *MUSK* | ATCACCACGCCTCTTGAAAC | TGTCTTCCACGCTCAGAATG |
| *AchRγ* | AGTGCAGGCAGTATTGGAGA | AGGTTACAGGCATCCACACAG |
| *MyoG* | TGAATGCAACTCCCACAGC | GCAACAGACATATCCTCCACC |
| *RUNX* | CGGCAGAACTGAGAAATGCT | CAACTTGTGGCGGATTTGTA |

**Table S2.** Antibodies used in this study

| *Antibody* | *Customer* | *Dilution* |
|---|---|---|
| *Goat Anti rabbit-HRP* | BIO-RAD 1706515 | 1:2000 WB |
| *Goat Anti mouse-HRP* | BIO-RAD 1706516 | 1:2000 WB |
| *MitoProfile® Total OXPHOS Rodent WB Antibody Cocktail* | MitoSciences MS604 | 1:1000 WB |
| *Mouse anti-GAPDH* | Abcam ab8245 | 1:10000 |
| *Mouse anti-ACOX1* | Santacruz-517306 | 1:1000 WB |
| *Rabbit anti-ACOX1* | Proteintech 10957-1-AP | 1:1000 WB |
| *Rabbit anti-LC3* | Sigma L7543 | 1:1000 WB |
| *Rabbit anti-Pex16* | 14816-1-AP | 1:1000 WB |
| *Rabbit anti-Pex5* | CST 83020S | 1:1000 WB |
| *Rabbit anti-PGC1-alpha+beta* | Ab54481 | 1:1000 WB |
| *Rabbit anti-FLNC* | Abclonal A13018 | 1:1000 WB |
| *Rabbit anti-CapZ* | 25043-1-AP | 1:1000 WB |
| *Rabbit anti-MYOZ* | 23880-1-AP | 1:1000 WB |
| *Rabbit anti-Alpha-Actinin* | Proteintech 24378-1-AP | 1:1000 WB |
| *Rabbit anti-PMP70* | Sigma P0497-200UL | 1:1000 WB; 1:100 IF |
| *Rabbit anti-VDAC* | CST 4866S | 1:1000 WB; 1:100 IF |
| *Rabbit anti-NCAM* | Sigma AB5032 | 1:100 IF |
| *Rabbit anti-Dystrophin* | Abcam ab15277 | 1:100 IF |
| *Rabbit anti-ACAA1* | Sigma HPA007244 | 1:100 IF |
| *Anti-VAMP1* | homemade[1] | 1:200 IF |

1. Rossetto, O. *et al.* VAMP/synaptobrevin isoforms 1 and 2 are widely and differentially expressed in nonneuronal tissues. *J Cell Biol* **132**, 167–179 (1996).
